# PRC2 direct transfer from G-quadruplex RNA to dsDNA: Implications for RNA-binding chromatin modifiers

**DOI:** 10.1101/2022.11.30.518601

**Authors:** Wayne O. Hemphill, Regan Fenske, Anne R. Gooding, Thomas R. Cech

**Author notes:** **Author contributions:** W.O.H. and T.R.C. designed research; W.O.H. and R. F. performed research; A.R.G. purified and validated proteins; W.O.H. and R.F. analyzed data; and W.O.H. and T.R.C. wrote the paper. **Competing Interest Statement:** T.R.C declares consulting status for Storm Therapeutics, Eikon Therapeutics, and SomaLogic. The authors have no other competing interests to declare.

## Abstract

The chromatin-modifying enzyme, Polycomb Repressive Complex 2 (PRC2), deposits the H3K27me3 epigenetic mark to negatively regulate expression at numerous target genes, and this activity has been implicated in embryonic development, cell differentiation, and various cancers. A biological role for RNA binding in regulating PRC2 histone methyltransferase activity is generally accepted, but the nature and mechanism of this relationship remains an area of active investigation. Notably, many *in vitro* studies demonstrate that RNA inhibits PRC2 activity on nucleosomes through mutually antagonistic binding, while some *in vivo* studies indicate that PRC2’s RNA-binding activity is critical for facilitating its biological function(s). Here we use biochemical, biophysical, and computational approaches to interrogate PRC2’s RNA and DNA binding kinetics. Our findings demonstrate that PRC2-polynucleotide dissociation rates are dependent on the concentration of free ligand, indicating the potential for direct transfer between ligands without a free-enzyme intermediate. Direct transfer explains the variation in dissociation kinetics reported previously, allows reconciliation of prior *in vitro* and *in vivo* studies, and expands the potential mechanisms of RNA-mediated PRC2 regulation. Moreover, simulations indicate that such a direct transfer mechanism could be obligatory for RNA to recruit proteins to chromatin.

**Significance:** Studies of PRC2 *in vitro* indicate that RNA inhibits its histone methyltransferase (HMTase) activity through mutually antagonistic binding with nucleosomes, but some *in vivo* studies paradoxically suggest that RNA binding is necessary to facilitate chromatin occupancy and HMTase activity. Our findings unveil a protein-intrinsic mechanism for directly exchanging RNA and DNA/nucleosome in PRC2’s binding site(s), which reconciles these prior findings by allowing antagonistic or synergistic RNA-mediated regulation dependent on RNA-nucleosome proximity. Furthermore, there is an increasing awareness that multiple chromatin-associated proteins exhibit regulatory RNA binding activity, and our findings indicate this “direct transfer” mechanism may be generally required for such regulation.

## Introduction

PRC2 is a histone methyltransferase (HMTase) that sequentially deposits three methyl groups onto lysine 27 of histone H3 (H3K27me1/2/3; (1–4), reviewed in refs (5–7)), and its activity is crucial for epigenetic silencing during development and cancer (5). How PRC2 is targeted to genetic loci is of considerable interest, given its critical function and abundance of target genes (8). PRC2’s core subunits include the EZH2 catalytic domain, SUZ12 scaffold subunit, EED histone tail-binding subunit, and RBBP4 histone chaperone subunit, and it has additional accessory subunits that define the PRC2.1 and PRC2.2 subtypes and differentially regulate its activity ((9–11), reviewed in ref (5)). PRC2 binds numerous long noncoding RNAs (lncRNAs) and pre-mRNAs in cell nuclei, and this RNA binding is believed to regulate PRC2’s HMTase activity (12, 13). Furthermore, biochemical studies have demonstrated that PRC2 has specificity for G-tracts and G-quadruplex (G4) RNA structures (14), which are ubiquitous in the human transcriptome, consistent with its widespread RNA binding in cells.

The nature and mechanism(s) of PRC2 regulation by RNA remain quite controversial. While some studies have proposed a role for RNA in PRC2 recruitment to chromatin (13, 15), others have suggested roles in PRC2 eviction from chromatin and/or inhibition of PRC2 catalytic activity (16–20), and these ideas are not mutually exclusive. Biochemical experiments have convincingly demonstrated that RNA antagonizes PRC2 HMTase activity (16, 18, 20), and that this is mediated by competitive binding with nucleosomes rather than catalytic suppression (17, 18). On the other hand, a recent work has demonstrated that the PRC2-RNA interaction is critical *in vivo* for maintaining H3K27me3 levels and chromatin occupancy at PRC2 target genes in induced pluripotent stem cells (21). It is prudent to note that the biochemical studies have utilized RNA and nucleosome in free solution, which is not representative of the chromatin-associated nascent RNA suspected to regulate PRC2 activity *in vivo* (16, 17, 19, 20). Furthermore, the role(s) of RNA in PRC2 activity could be contextual to chromatin architecture, available PRC2 accessory subunits and protein partners, competing RNA-binding proteins, and/or post-translational modifications. Thus, prior biochemical studies may lack considerations relevant to *in vivo* function. Direct evidence for a mechanism that can reconcile RNA antagonizing PRC2’s nucleosome binding and HMTase activity *in vitro* with RNA-mediated PRC2 recruitment *in vivo* has yet to be reported.

Herein, we measure the kinetics of PRC2’s RNA and DNA binding using biochemical, biophysical, and computational methods. Our findings unexpectedly reveal that PRC2 has the intrinsic ability to exchange one nucleic acid for another without completely dissociating from the first nucleic acid. Such mechanisms have been well-studied for homo-multimeric DNA-binding proteins like lac repressor (22), *E. coli* catabolite activator protein (CAP) (23), SSB (24), and recA (25) and for the hexameric RNA-binding protein Hfq (26). Historically, this phenomenon has been variously identified as “concentration-dependent dissociation”, “direct transfer” (23–25), “facilitated exchange” (27), or “active exchange” (26), and it is related to the sister phenomena of protein movement along DNA by looping (28, 29) and “facilitated dissociation” (30). The proteins in these cases are homo-oligomeric, and as others have noted (31), their multiple ligand-binding sites likely facilitate direct transfer by providing a foothold for a second ligand before it displaces a previously bound ligand. We propose that PRC2’s ability to directly transfer from one nucleic acid to another, despite lacking the structural symmetry of prior proteins exhibiting direct transfer, may reconcile the disparate eviction-versus-recruitment models of previous studies. Furthermore, binding to nascent RNA has been suggested as a general strategy by which transcription factors and other DNA-binding proteins are maintained at high local concentrations. Our findings indicate that this model may be feasible only if the protein can directly transfer from RNA to DNA without dissociation, suggesting direct transfer capabilities may have general relevance.

## Results

### PRC2’s Dissociation Rate from G4 RNA is Competitor Concentration-Dependent

Two prior studies from our group (17, 21) used fluorescence polarization-based competitive dissociation (FPCD) experiments to determine the dissociation rate constant for a G4 RNA species from PRC2, but they obtained significantly different values despite nearly identical methodologies. One of the few methodological distinctions between these two studies was the concentration of unlabeled competitor RNA (decoy). Long et al. and Wang et al. used a 200- and 2000-fold excess of decoy over fluorescently labeled RNA (prey), respectively, both of which should have been sufficient to totally prevent prey rebinding. We replicated these experiments across a range of decoy concentrations (Fig. 1 and Supp. Fig. 1) to ensure a sufficient excess of decoy was achieved. Unexpectedly, the observed dissociation rate of PRC2 and G4 RNA did not plateau at excess concentrations of decoy, but instead continued to increase linearly in a decoy concentration-dependent manner (Fig. 1d). Such behavior can be explained by the incoming competitor (decoy) displacing the initially bound molecule (prey) without free protein as an intermediate (i.e., direct transfer of the protein between the two ligand molecules) (24). Under such a model, the discordance between the original Wang et al. and Long et al. apparent dissociation rate constants is expected, and both findings are consistent with our data given the respective decoy concentrations used.

**Fig 1.**
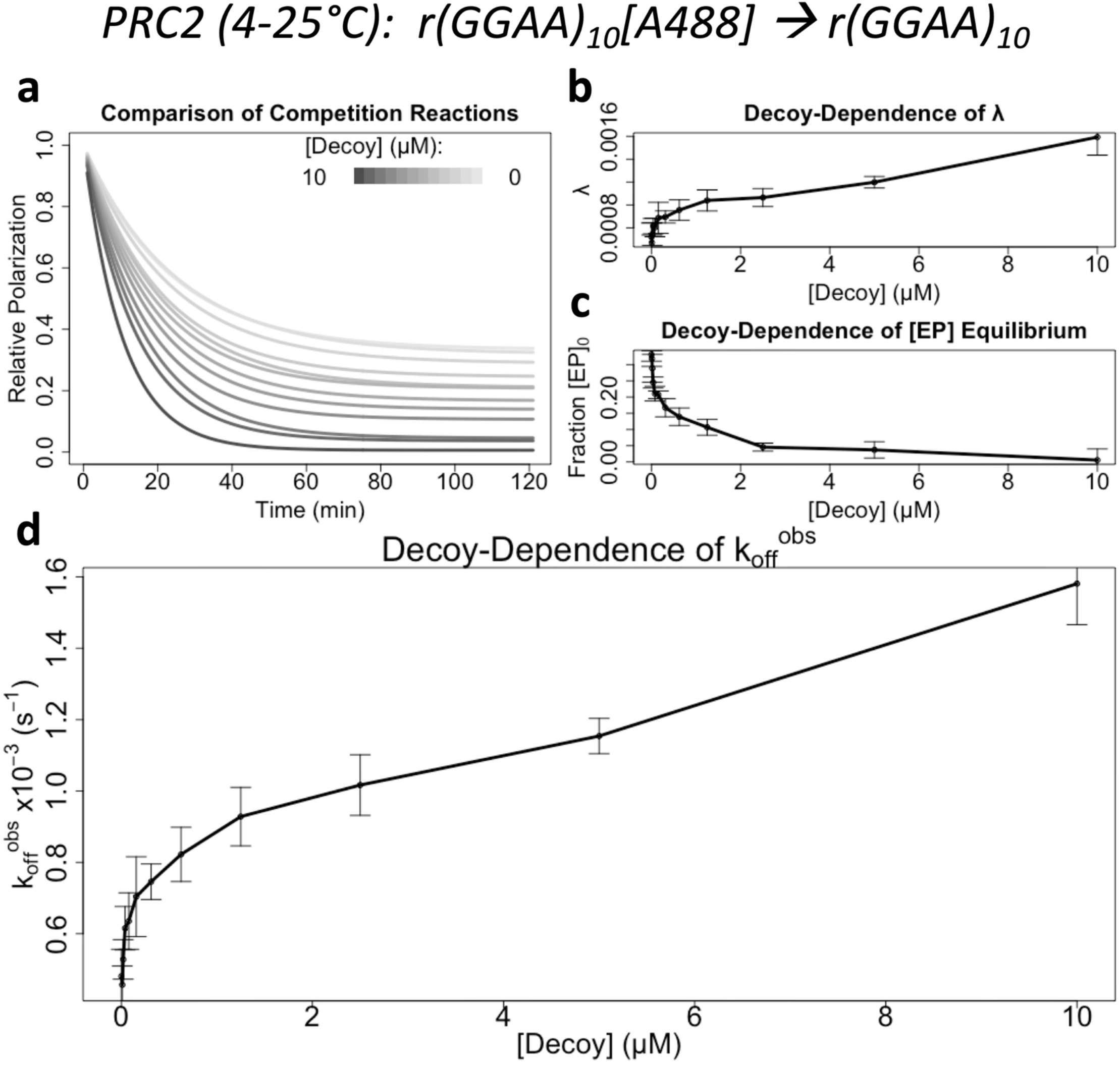
PRC2’s Observed Dissociation Rate from a G4 RNA is Dependent on Competitor Concentration. Fluorescence polarization-based competitive dissociation experiments (Fig. 2) were performed as described to replicate the original Wang et al and Long et al experiments over a range of competitor (‘decoy’) RNA concentrations. Data are from a single representative experiment (of n=3), where error bars indicate mean ± SD for four technical replicates. **[a]** Reaction data for each competitor concentration was regressed with an exponential dissociation equation, and the fit lines from each condition are shown for comparison. **[b]** The regressions’ decay constant values (λ, see Eq. 3.1) plotted versus competitor concentration. **[c]** The reactions’ proportions of initial complex remaining at equilibrium ([EP]_0_, see Eq. 3.1 – N_min_) plotted versus competitor concentration. **[d]** The reactions’ observed initial dissociation rates (k_off_^obs^, see Eq. 3.2) plotted versus competitor concentration. Solid lines in panels b-d are visual aids connecting data means.

### PRC2 Exhibits Direct Transfer between G4 RNA and dsDNA

To validate the above result, we performed additional FPCD experiments and analyzed their data with regression models that exclude or allow such direct transfer (Fig. 2). First, since the initial experiments utilized an RNA with 10 G-tracts that could form G4s heterogeneously, we tested a simpler RNA sequence containing only four G-tracts and found that it also exhibited direct transfer kinetics (Supp. Fig. 2a and Table 1). Next, since prior competition reactions were initially incubated at 4°C to slow off-rates but warmed to 25°C in the plate reader, we repeated this experiment at constant room temperature (Supp. Fig. 2b and Table 1) to interrogate temperature-dependent effects. Then, we repeated the experiment using a carrier poly(A) RNA that does not bind PRC2 to keep total RNA concentration constant in the reactions (Supp. Fig. 2c and Table 1), so that any nonspecific polynucleotide concentration-dependent phenomena (e.g., electrostatic effects) could be ruled out as artifactual explanations. Finally, we repeated the experiment with a different fluorescent label on the prey molecule (Supp. Fig. 2d and Table 1) to interrogate interactions with the fluorophore. The data from all three validation experiments were well fit by a regression model allowing direct transfer kinetics but poorly fit by a classic model of competition, and thus favored PRC2 intrinsic direct transfer between G4 RNA molecules.

**Table 1.**
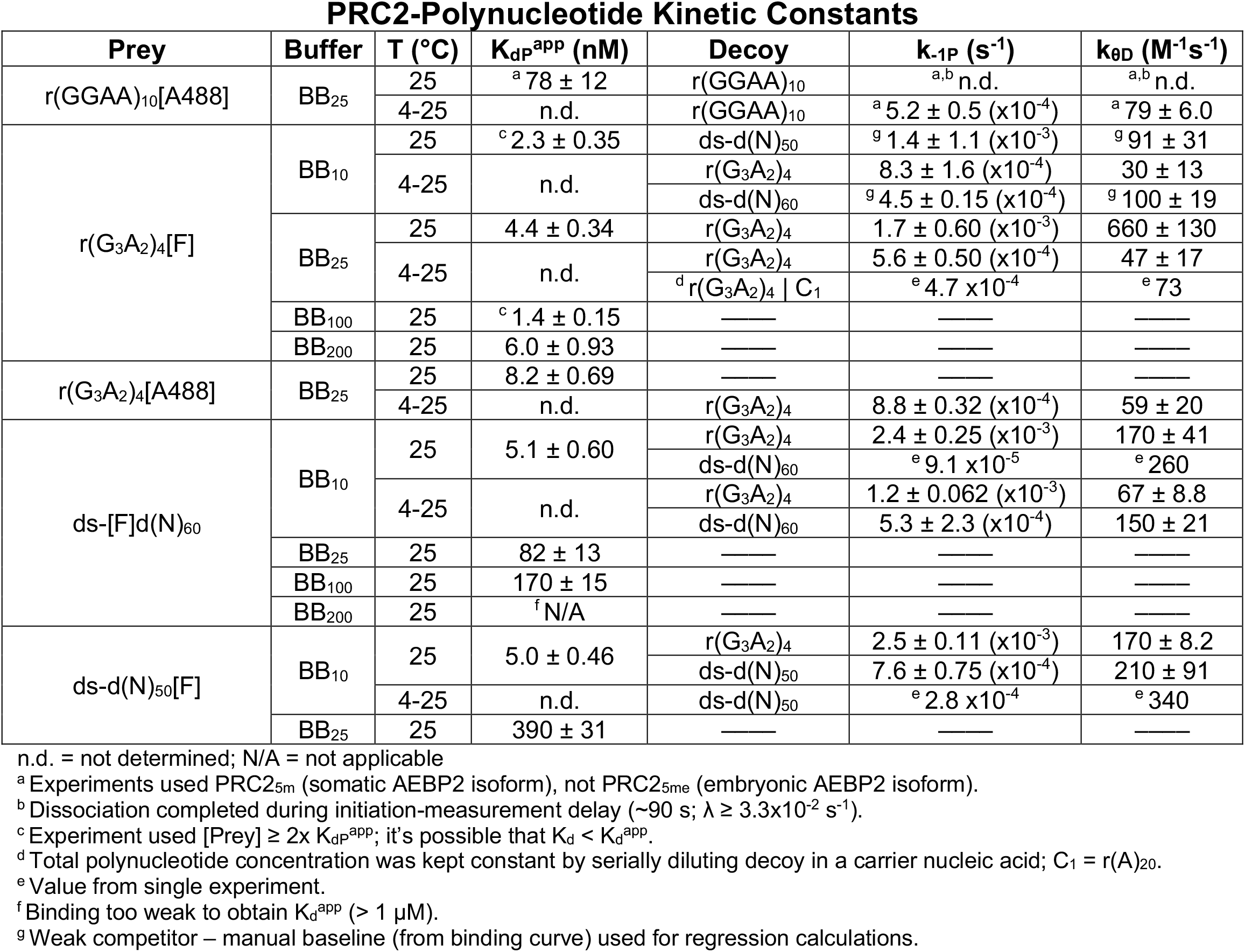
Rate Constants for a Variety of Protein-Ligand Interactions. Fluorescence polarization-based methodology (Supp. Fig. 1) was used as described to determine the apparent equilibrium dissociation constants (K_dP_^app^), intrinsic dissociation rate constants (k_-1P_), and direct transfer rate constants (k_θD_) for several protein-ligand interactions. Values indicate mean ± SD for at least three independent experiments.

**Fig 2.**
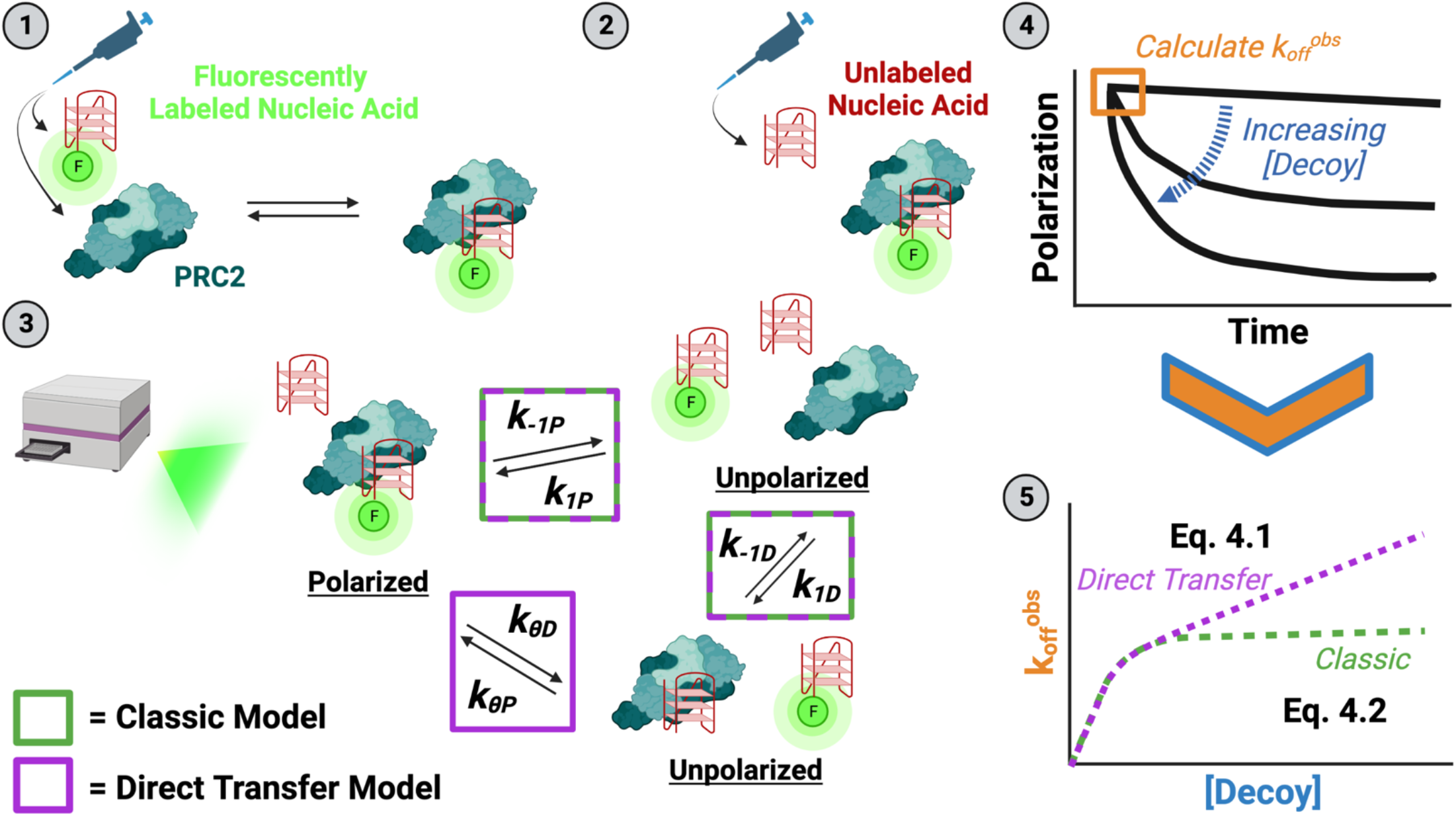
Experiment and Analysis Strategy to Measure Direct Transfer Kinetics. (1) The minimum amount of PRC2 required for saturated binding is mixed with a trace amount of fluorescently labeled nucleic acid (prey), then incubated (at 4/25 °C) until thermal and reaction equilibrium. (2) Various concentrations of unlabeled nucleic acid (decoy) are added to the preformed complex to initiate reactions (at 25°C). (3) The time-course reactions are immediately monitored by fluorescence polarization in a microplate reader (at 25°C). Potential complexes with their polarization states are shown, and they are labeled with rate constants describing inter-complex transitions. Rate constants associated with a classic competition model are indicated by green boxes, and those additionally necessary for a direct transfer model are indicated by a purple box. The inter-complex transition solely associated with the direct transfer model has an implied unstable ternary complex intermediate. The system of differential equations describing these reactions is given by Eq. 1. (4) Polarization signals are normalized to the range in polarization signal across all decoy concentrations to give proportion of initial complex remaining. Normalized polarization signals are plotted versus time and fit with one-phase exponential decay regression. (5) The regressions’ initial slopes (k_off_^obs^) are plotted versus decoy concentration and regressed with custom equations describing the classic competition and direct transfer models to determine rate constant values.

Of particular biological relevance is PRC2’s potential for direct transfer between RNA and chromatin. Prior studies indicate PRC2’s affinity for nucleosomes is almost entirely mediated by exposed nucleosome linker DNA (17), suggesting comparable-length dsDNA species should be representative of PRC2’s nucleosome binding activity. Thus, we performed FPCD experiments with our simple G4 RNA and a 60-bp dsDNA, using all possible prey-decoy combinations (Fig. 3). Notably, our results indicate that direct transfer occurs between all species. However, they indicate that RNA→dsDNA direct transfer is more efficient (k_θD_/k_-1P_, see Fig. 2) than dsDNA→RNA direct transfer (Table 1). The relevance of our metric for direct transfer efficiency is supported by our concurrent studies (companion manuscript). Similar experiments with a 50-bp dsDNA species echoed these findings (Supp. Fig. 3). We also note that prior reports of PRC2 dsDNA and G4 RNA binding affinities (13, 14, 17, 21) are consistent with our corresponding values in Table 1.

**Fig 3.**
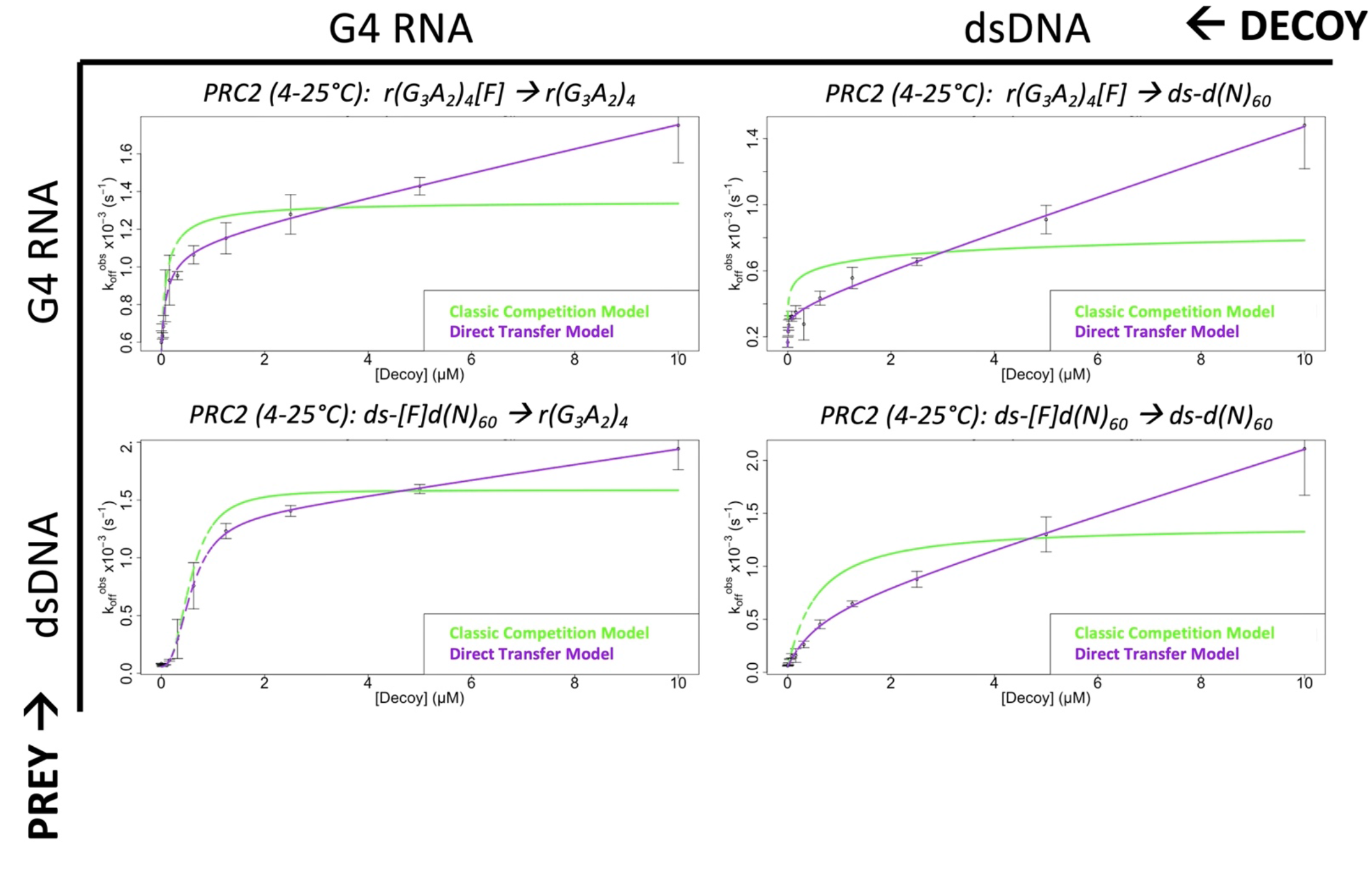
PRC2 Exhibits Direct Transfer Kinetics for G4 RNA and dsDNA. FPCD experiments (Fig. 2) were performed (buffer = BB_10_) as described for every prey-decoy combination of a G4 RNA and 60-bp dsDNA. Data are from representative experiments (of n ≥ 3), where error bars indicate mean ± SD for four technical replicates.

### PRC2 Makes Additional Electrostatic Contacts with dsDNA Not Utilized for G4 RNA

Prior studies indicate G4 RNA and dsDNA binding to PRC2 are mutually antagonistic (i.e., competitive) (17, 18), which suggests that our observed direct transfer events for PRC2 ligands involve competition for shared protein-polynucleotide contacts. Yet, the direct transfer kinetics for RNA→dsDNA events are more efficient (defined above) than for RNA→RNA or dsDNA→RNA events, implicating additional protein-polynucleotide contacts unique to one or both ligands. To interrogate this possibility, we used FP to determine K_d_^app^for G4 RNA and dsDNA at a range of salt concentrations. The experiments demonstrated a significantly greater influence of ionic strength on PRC2’s binding affinity for dsDNA than for G4 RNA (Fig. 4a). More specifically, linear regression of log(K_d_^app^) vs log([KCl]) plots (Fig. 4b) indicates that more salt bridges mediate PRC2 binding to dsDNA (m ≈ 1.4 ± 0.68) versus G4 RNA (m ≈ 0 ± 0.34) (32). This is consistent with the previous conclusion that the PRC2-RNA interaction is not primarily electrostatic (33), though a different RNA species was used. These data suggest that PRC2 has additional ionic contacts with dsDNA that are not utilized during its binding to G4 RNA.

**Fig 4.**
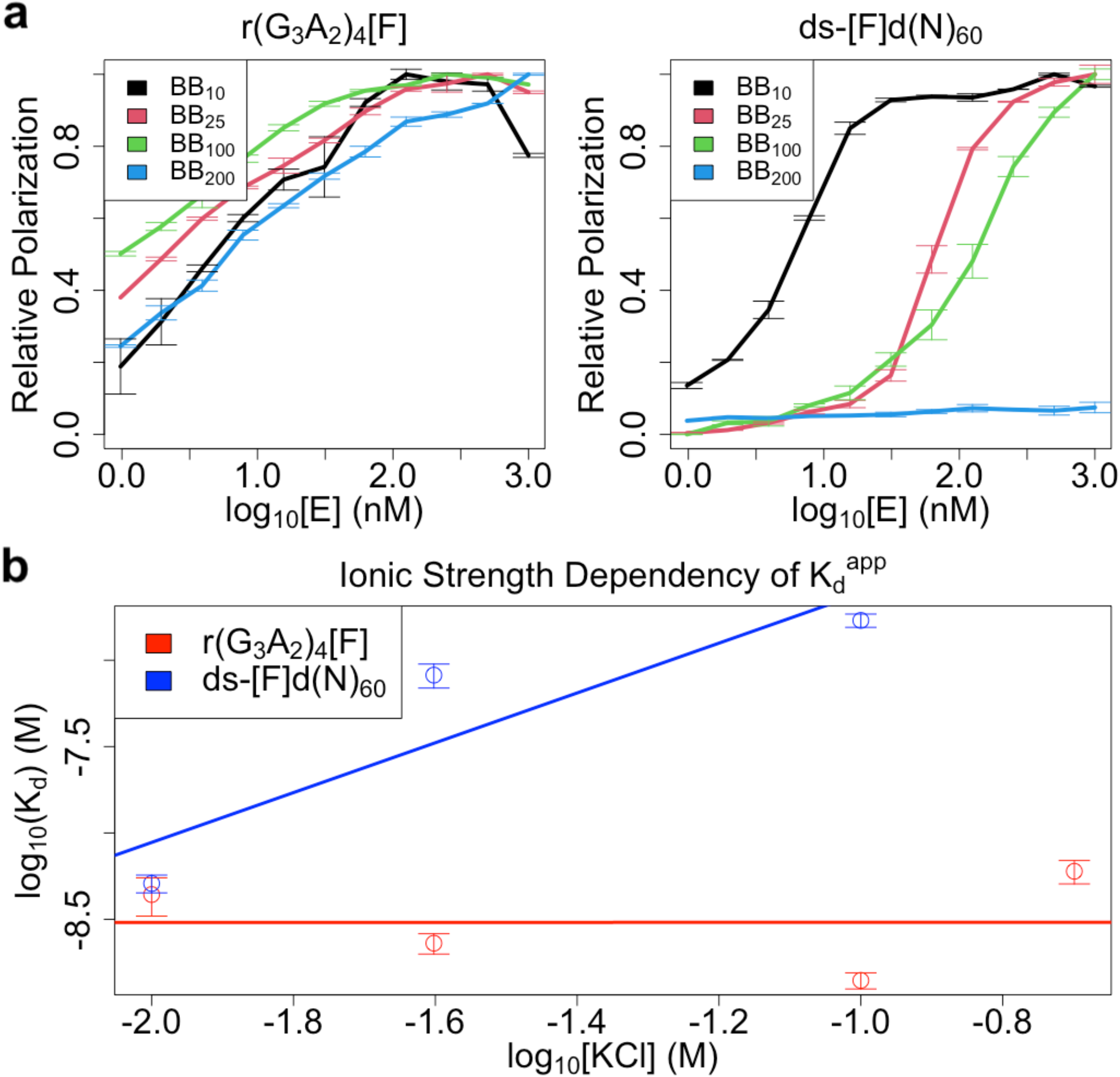
Ionic Interactions Contribute to PRC2’s dsDNA but not G4 RNA Affinities. FP-based equilibrium binding curve experiments were carried out under various salt concentrations (BB_X_ = X mM KCl) for a G-quad RNA and 60-bp dsDNA, then salt-dependence of their K_d_^app^ compared. **[a]** Binding curves for indicated PRC2 ligands. Curves are composites of all replicates/experiments, where error bars indicate mean ± SD. **[b]** Affinity versus ionic strength plots with K_d_ ^app^ from regression of data in panel a. Data points are composites of all experiments in panel a, where error bars indicate mean ± SD.

### Modeling Suggests the PRC2 Direct Transfer Mechanism Allows RNA-Mediated Recruitment to Nucleosomes

Prior studies indicate that RNA inhibits PRC2’s nucleosome binding and HMTase activity (13, 17), while others paradoxically suggest that RNA facilitates PRC2 chromatin occupancy and H3K27me3 deposition (21). To interrogate whether PRC2 direct transfer might reconcile these views, we constructed a reaction scheme representative of PRC2’s proposed biochemical activity (Fig. 5a). This scheme accounts for classic PRC2 (E) binding to (k_1_) and dissociation from (k_-1_) RNA (R) and nucleosomes (N), PRC2 catalytic (k_cat_) methylation of nucleosomes (N^m^), and the potential *in vivo* proximity (α) between nascent RNA and chromatin that was not recapitulated by prior *in vitro* studies. We simulated reactions under this scheme using our empirically determined rate constants for association, dissociation, and direct transfer events (Table 1). The results (Fig. 5b and Supp. Fig. 4) indicate that RNA should be antagonistic to PRC2 HMTase activity in free solution (α = 1), as observed experimentally. However, RNA should become synergistic as RNA-nucleosome proximity increases (α > 1).

**Fig 5.**
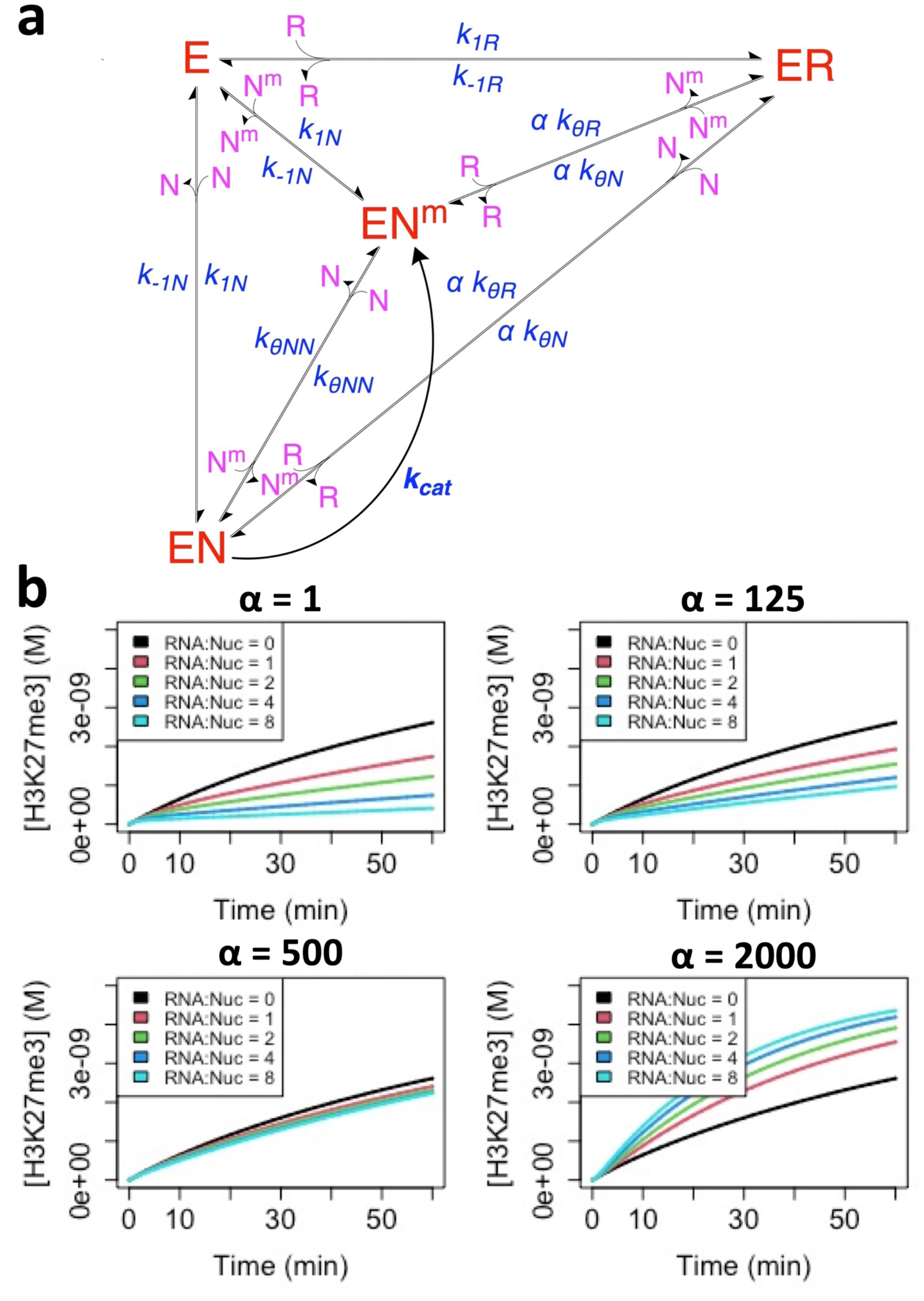
Direct Transfer Allows RNA to Boost PRC2 HMTase Activity. **[a]** Reaction scheme includes PRC2 (E), RNA (R), nucleosomes (N), and methylated nucleosomes (N^m^), where conjugates are complexes of the respective reactants. Major PRC2 states are shown in red, additional reactants in purple, and rate constants in blue. For rate constants, k_1_ is for association, k_-1_ for dissociation, k_θ_ for direct transfer, k_cat_ for catalysis, and α for effective molarity changes from reactant proximity (see Eq. 5). Inter-complex transitions defined by the k_θ_ rate constants are like those shown in Fig. 2, and their removal collapses this scheme to a classic model. The α parameter is included for any PRC2 complex transitions where RNA-nucleosome direct transfer is possible. **[b]** Empirically determined rate constants (Table 1) were used to simulate PRC2 HMTase activity for a range of effective molarities (α) and RNA-nucleosome molar ratios (RNA:Nuc). Black curves represent HMTase time-course reactions in the absence of RNA, and the colored lines represent the effect of increasing RNA concentrations.

As expected, all RNA effects on PRC2 HMTase activity were ablated if the PRC2-RNA complex was unstable (Supp. Fig. 5). Notably, the ability of RNA to boost PRC2 HMTase activity under high RNA-nucleosome proximity is completely dependent on direct transfer (Supp. Fig. 6). Overall, these data demonstrate that PRC2’s mutually antagonistic RNA and nucleosome binding are reconcilable with RNA-mediated recruitment of PRC2 to target genes, but only if PRC2 can direct transfer from RNA to nucleosomes.

### Direct Transfer May Be Generally Required for RNA Recruitment of Chromatin Proteins

In the case of PRC2 and some other chromatin-associated proteins, RNA and nucleosomes bind mutually antagonistically (competitively). However, other proteins can stably bind both RNA and chromatin simultaneously. For example, the transcription factor Yin Yang 1 (YY1) binds DNA and RNA independently, and Sigova et al (34) proposed that its RNA binding keeps YY1 trapped near its DNA binding sites to help recruit it to chromatin DNA. To interrogate this alternative situation of simultaneous binding, we designed a reaction scheme for a hypothetical HMTase enzyme with independent RNA and nucleosome binding activity (Fig. 6a). This allows direct comparison of the behavior of the mutually antagonistic versus independent binding situations. This scheme accounts for effective molarity (α) changes due to nucleosome-RNA proximity, RNA-mediated suppression of catalysis (β), and RNA/nucleosome effects on nucleosome/RNA binding affinity (δ). We then simulated reactions with a range of values for these parameters, using kinetic constants consistent with those reported by Sigova et al. for YY1 (34). As expected, our results (Fig. 6b) support RNA concentration having no effect on activity in free solution (α = 1), assuming that RNA binding does not affect catalysis (i.e., β = 1). In contrast, the simulations show that RNA concentration facilitates catalytic activity as RNA-nucleosome proximity increases (α > 1). However, we note that this synergy is sensitive to relative reactant concentrations, and it is easily ablated by even minor RNA-mediated catalytic suppression (β < 1) (Supp. Fig. 7).

**Fig 6.**
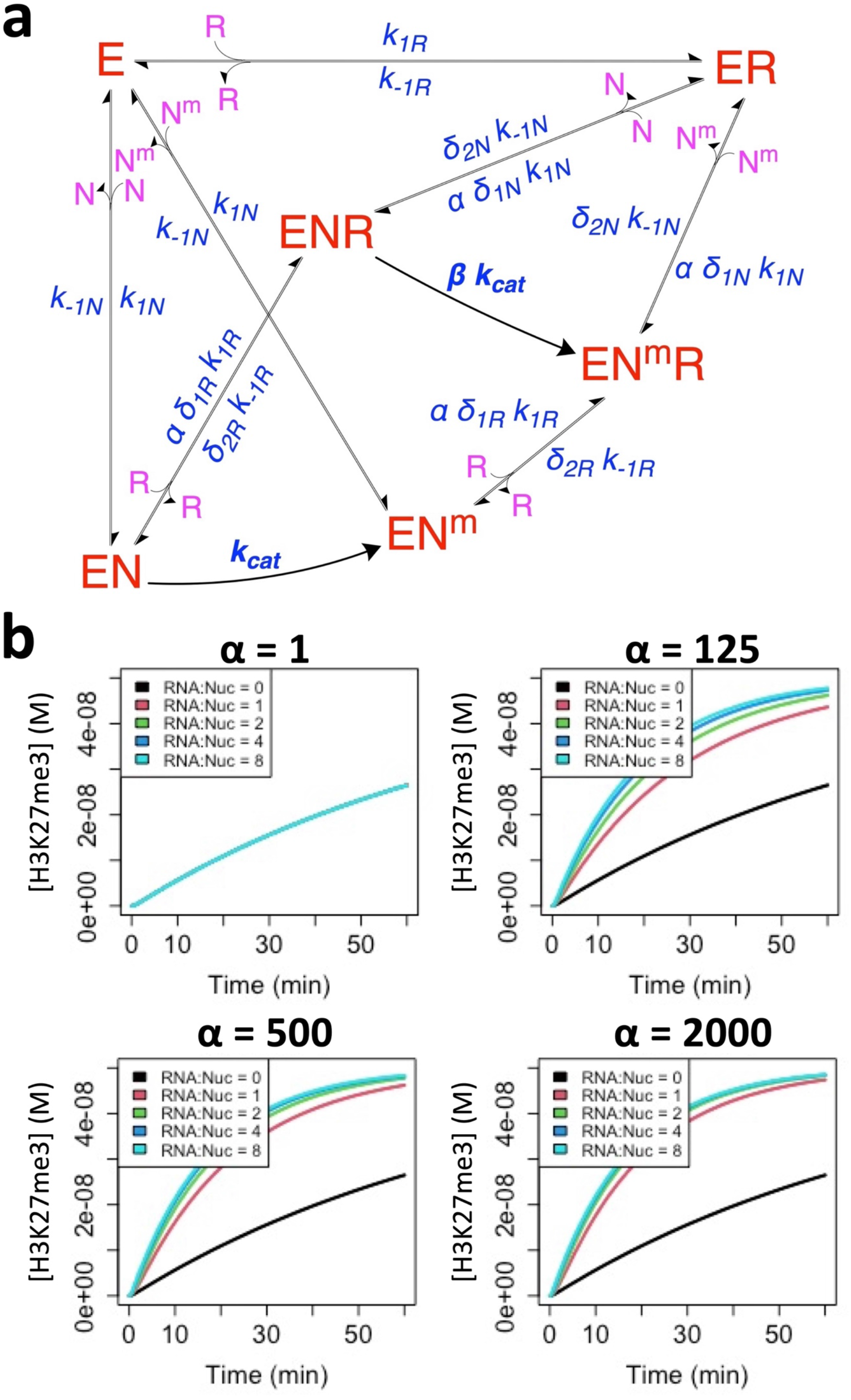
Stable RNA and Nucleosome Co-Binding Could Boost a Protein’s Activity. **[a]** Reaction scheme for protein with independent RNA and nucleosome binding includes protein (E), RNA (R), nucleosomes (N), and methylated nucleosomes (N^m^), where conjugates are complexes of the respective reactants. Major protein states are shown in red, additional reactants in purple, and rate constants in blue. Regarding rate constants, k_1_ is association, k_-1_ is dissociation, k_cat_ is catalysis, α is effective molarity changes from reactant proximity (see Eq. 5), β is effect of bound RNA on catalytic activity, and δ is effect of bound RNA/Nucleosome on ternary Nucleosome/RNA binding. **[b]** The Sigova et al kinetic constants for YY1 were used to simulate catalytic activity for a range of effective molarities (α) and RNA-nucleosome molar ratios (RNA:Nuc). Black curves represent HMTase time-course reactions in the absence of RNA, and the colored lines represent the effect of increasing RNA concentrations.

Importantly, this synergy is completely dependent on formation of the stable ternary complex, and preventing its formation creates an antagonistic relationship between RNA and activity (Supp. Fig. 8). Thus, direct translocation between RNA and chromatin DNA without a free-enzyme intermediate is necessary to improve catalytic activity for the alternative situation of noncompetitive binding. These data address chromatin binders with catalytic activity, but recent findings (35) suggest that RNA-binding transcription factors (RBTFs) like YY1, whose biological activity is not catalytic in nature, may be quite prevalent. To interrogate the relevance of our findings to such RBTFs, we eliminated catalytic activity from the prior reaction scheme (Fig. 6a) and alternatively monitored nucleosome binding in simulations. Our results suggest RNA could improve RBTF activity at high RNA-nucleosome proximity without any detrimental effects in free solution (Supp. Fig. 9a); this positive effect of RNA binding is dependent on an RBTF’s ability to function on their nucleosome target with RNA co-bound (Supp. Fig. 9b) and on formation of the stable ternary complex (Supp. Fig. 9c). Consequently, translocation without free enzyme intermediates seems necessary for RNA-mediated facilitation of activity for proteins with independent RNA and nucleosome binding, independent of whether the protein acts catalytically or simply by binding DNA.

These findings indicate that PRC2-like systems and independent binding (YY1-like) systems can both have RNA-mediated facilitation of their activity, and that this facilitation is dependent on the ability to translocate between RNA and nucleosomes without a free-protein intermediate. While PRC2 would accomplish this through direct transfer, independent binding systems accomplish this through a stable ternary complex. However, while our PRC2 reaction scheme for direct transfer events (Fig. 5a) doesn’t explicitly identify a ternary complex like the scheme for independent binders (Fig. 6a), the existence of an unstable ternary complex intermediate is still implied. Indeed, making the ternary complex for an HMTase with independent binding (Fig. 6a) 100-fold less stable (i.e., δ_2_ = 100) produces a similar relationship between activity, RNA concentration, and RNA-nucleosome proximity as a PRC2-like system (Supp. Fig. 10). We note a comparable trend for independent binders without catalytic activity, like RBTFs (Supp. Fig. 9d). Collectively these data suggest that the seemingly distinct PRC2-like and Sigova et al. models for RNA-mediated recruitment to chromatin both rely on translocation between RNA and nucleosome DNA through a ternary complex intermediate, and they differ only in the stability of their ternary complexes. Thus, YY1-like systems rely on the same mechanics intrinsic to direct transfer, but they exhibit different kinetics for the mechanism (i.e., the lifetime of the ternary intermediate). Thus, some form of direct transfer may be generally necessary for RNA-binding chromatin-associated proteins to have their functions on chromatin facilitated by RNA.

In the above cases, our data indicate that direct transfer creates a synergistic relationship between RNA concentration and protein activity, but only if RNA-nucleosome proximity is high, suggesting direct transfer alone is insufficient for RNA-mediated facilitation of protein function. However, since our reaction schemes were incapable of accounting for the effects of RNA-nucleosome proximity without ternary complex formation or direct transfer, our findings don’t address whether RNA-nucleosome proximity alone is sufficient for RNA-mediated facilitation of protein function. To interrogate this question, we employed single-molecule dynamics simulations (SMD) of mutually exclusive protein binding to RNA and nucleosomes tethered together. Our findings indicate that increasing RNA binding affinity increases RNA occupancy (Fig. 7a) as expected, but slightly decreases nucleosome occupancy (Fig. 7b). Similarly, the increased intermolecular distance between nucleosomes and nearby unbound protein (Fig. 7c), and the reduced concentration of protein in nucleosome-adjacent solvent space (Fig. 7d), confirm an antagonistic relationship between RNA and protein function (nucleosome binding). Thus, RNA-nucleosome proximity alone is not sufficient for RNA-mediated recruitment to chromatin but is instead required alongside direct transfer.

**Fig 7.**
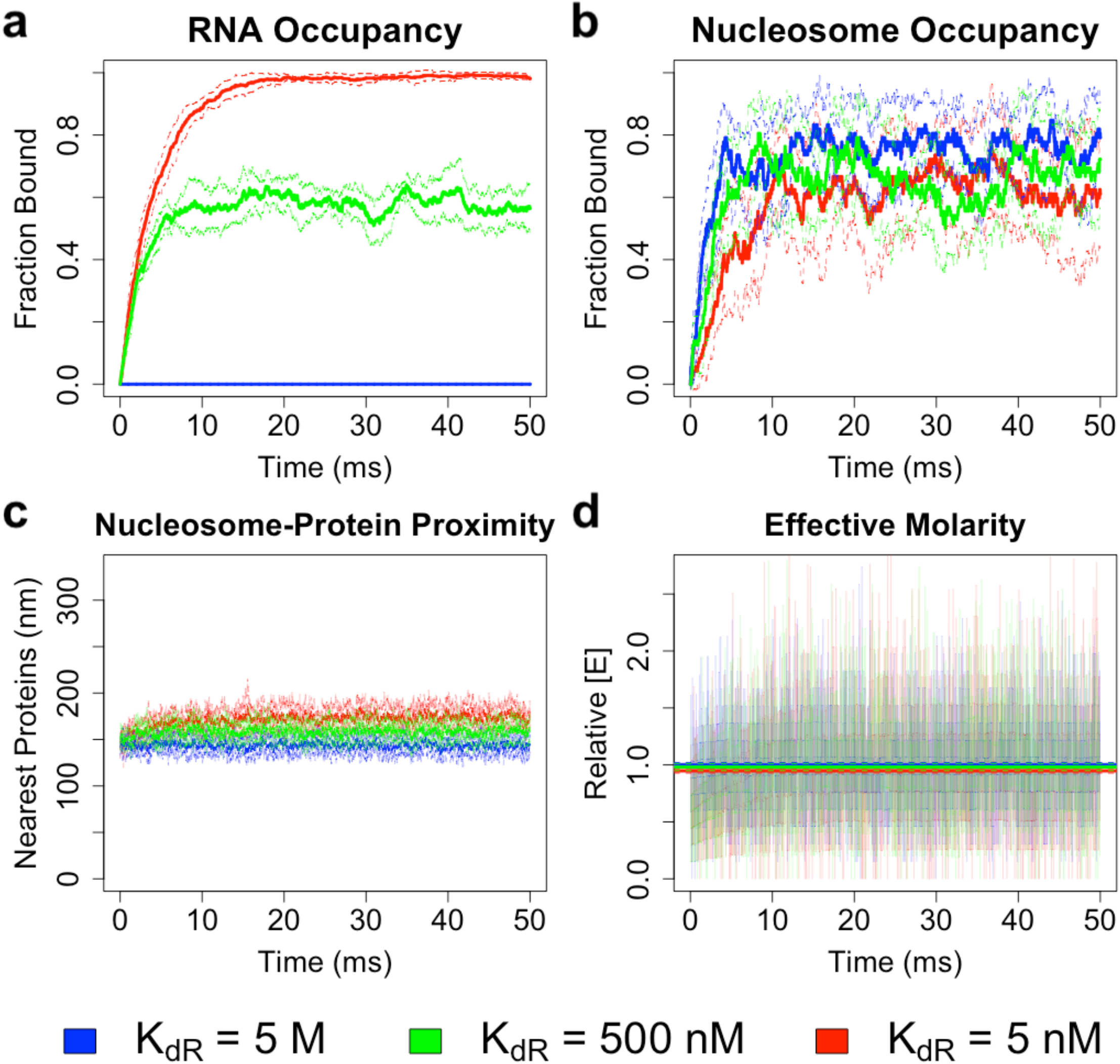
RNA-Nucleosome Proximity Alone Isn’t Sufficient to Improve Nucleosome Occupancy for Mutually Exclusive Binders. Single-molecule dynamics simulations were used to predict mutually exclusive protein binding to nucleosomes and tethered RNA (1:8 molar ratio), on the millisecond timescale, for a variety of RNA binding affinities. Colors correspond to different RNA binding affinities. **[a]** *Protein Association with RNA Over Time*. Solid and dashed lines indicate mean ± SD (across n=8 simulations), respectively, of the fraction of total RNA molecules bound. **[b]** *Protein Association with Nucleosomes Over Time*. Solid and dashed lines indicate mean ± SD (across n=8 simulations), respectively, of the fraction of total nucleosome molecules bound. **[c]** *Proximity of Nucleosomes to Nearest Protein Molecules Over Time*. For every simulation, the average intermolecular distances between each nucleosome molecule and its four closest protein molecules were calculated and averaged at every time point. Solid and dashed lines indicate mean ± SD (across n=8 simulations), respectively, of this average intermolecular distance. **[d]** *Effective Molarity Over Time*. For every simulation, the average concentration of unbound protein surrounding nucleosomes was divided by the concentration of unbound protein in the reaction to calculate relative effective molarity at every time point. Transparent lines indicate mean (across n=8 simulations) of effective molarity over time. For each simulation, the effective molarity was averaged over time, and solid and dashed lines indicate mean ± SD (across n=8 simulations), respectively, in average effective molarity.

## Discussion

### Implications for PRC2 Biology

Our biophysical studies indicate that PRC2 is capable of intrinsic direct transfer between G4 RNA and nucleosome-linker-sized dsDNA (Fig. 3 & Table 1), and our computational investigations reveal that this behavior should allow RNA to have either an antagonistic or a synergistic effect on PRC2 activity depending on RNA-nucleosome proximity (Fig. 5). These findings provide direct evidence for a mechanism (Fig. 8a) that allows RNA to facilitate PRC2’s HMTase activity, which can reconcile prior perplexing *in vitro* and *in vivo* results where RNA was alternatively found to inhibit PRC2 or recruit it to sites of action. We propose that PRC2 binding to nascent RNA (Fig. 8a – 1-2) could increase the effective molarity for direct transfer events (Fig. 8a – 3), allowing increased chromatin association and H3K27me3 deposition (Fig. 8a – 4) relative to an RNA-free (or RNA binding-free) system. It’s prudent to note that while these findings demonstrate that PRC2’s direct transfer kinetics can support RNA-mediated recruitment to chromatin, they do not prove its occurrence *in vivo*. Ideally, one would test *in vivo* a separation-of-function mutant that prevented direct transfer but retained full RNA and chromatin binding activities. However, we are pessimistic that such a mutant could be obtained, given that direct transfer is likely an intrinsic property of the nucleic acid-binding surfaces of PRC2.

**Fig 8.**
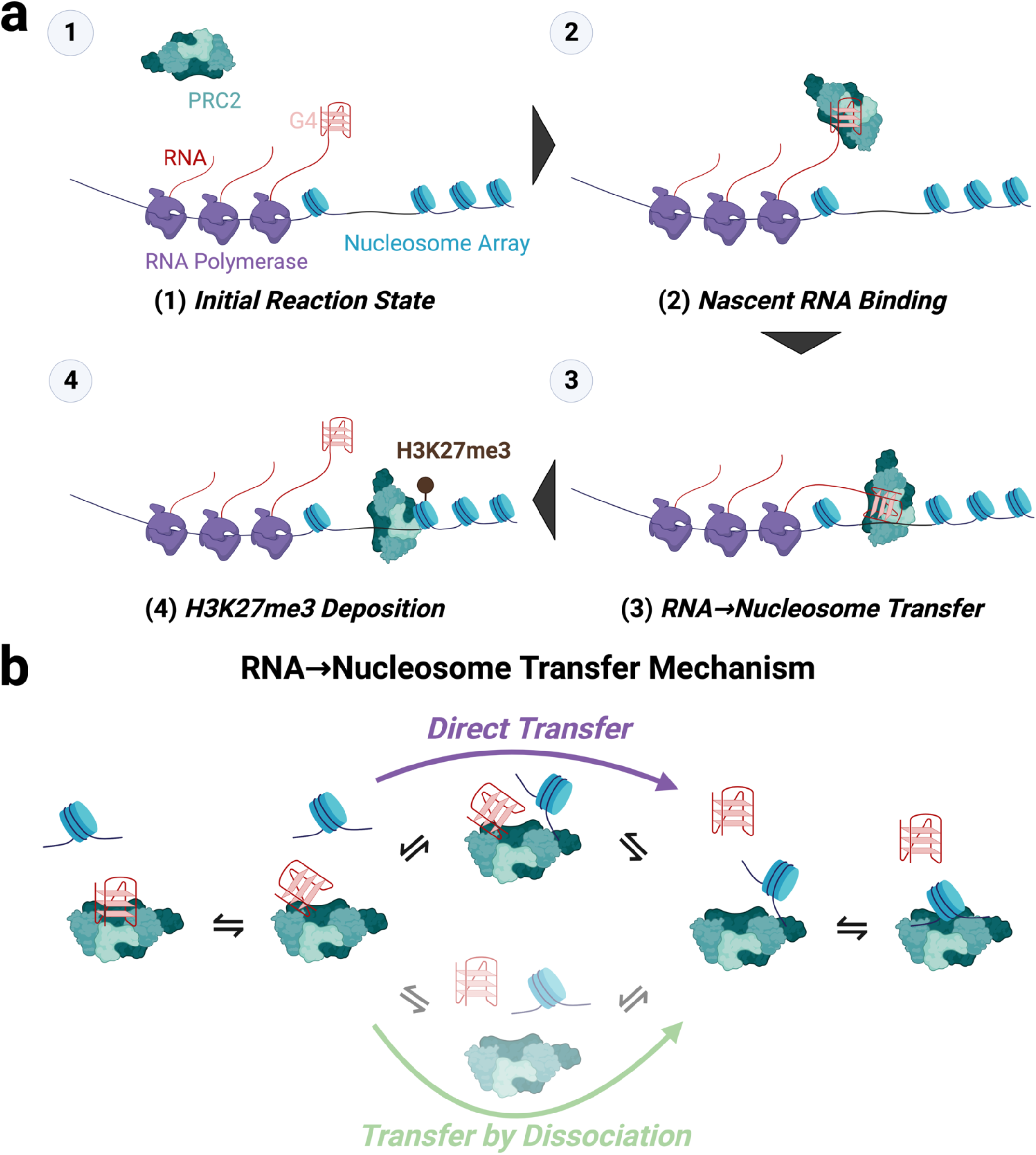
A Direct Transfer Model of RNA Regulation of PRC2 HMTase Activity. **[a]** *Proposed Steps for RNA Recruitment of PRC2 HMTase Activity*. (1) G4-containing nascent RNA at transcriptionally active PRC2 target genes (2) is bound by PRC2, (3) RNA-tethered PRC2 is transferred onto spatially proximal nucleosomes, then (4) PRC2 deposits its H3K27me3 mark. **[b]** *Proposed Mechanism for the Direct Transfer Step*. PRC2-bound G4 RNA (left) can transiently fluctuate to a partially bound state (middle left). From this state, the RNA can fully dissociate (middle bottom) before a nucleosome binds PRC2 (middle right & right) (Transfer by Dissociation), or a nucleosome can partially bind to create an unstable ternary complex (middle top) that elicits RNA dissociation (middle right) and full nucleosome association (middle right) (Direct Transfer).

The alternative situation, where RNA binding inhibits PRC2 activity, appears to occur *in vivo*. For example, it explains why many active genes have PRC2 close enough to be captured by ChIP (chromatin immunoprecipitation), yet the PRC2 does not act there (33). Furthermore, RNA inhibition of PRC2 has been shown in cells by Jenner et al (18, 19). Consistent with these observations, our simulations (Fig. 5) indicate that RNA is indeed antagonistic below a certain threshold of RNA-nucleosome proximity, although we have no way of predicting when and where these threshold conditions would be met *in vivo*. Importantly, our demonstration of PRC2’s direct transfer kinetics means that the recruitment and eviction models need not be mutually exclusive. As such, it’s possible that RNA-mediated regulation of PRC2 operates as a “switch,” where predominantly free versus predominantly nascent RNA landscapes around target genes drive PRC2’s relationship with RNA being antagonistic versus synergistic, respectively.

### Biophysical Mechanism for Direct Transfer

These findings demonstrate that PRC2 translocates directly between relevant polynucleotide species, but they raise compelling questions about the underlying biophysical mechanism that allows this direct transfer. Our data (Fig. 3 and Table 1) clearly indicate direct transfer occurs between nearly identical polynucleotides (labeled vs unlabeled), and between G4 RNA and dsDNA, which prior studies (17, 18) indicate have mutually antagonistic PRC2 binding. Collectively, these data implicate shared protein-polynucleotide interactions between competing ligands in a direct transfer mechanism. We propose a model (Fig. 8b) where PRC2-bound ligands can fluctuate to partially associated intermediates (Fig. 8b – left and middle-left), such that competitor ligands can occupy transiently exposed shared protein contacts to form an unstable ternary complex (Fig. 8b – top-middle); this prompts full dissociation of one ligand (Fig. 8b – middle-right) and allows full association of another ligand (Fig. 8b – right). This model is also consistent with our other findings, namely our salt dependency data (Fig. 4) indicating that dsDNA has additional ionic contact(s) with PRC2 not present for G4 RNA, and our kinetic data (Fig. 3 and Table 1) demonstrating variable relative efficiency among the dsDNA→G4 (low; k_θD_/k_-1P_ ≈ 5.6 ×10^4^ M^-1^), G4→G4 (moderate; k_θD_/k_-1P_ ≈ 8.3 ×10^4^ M^-1^), and G4→dsDNA (high; k_θD_/k_-1P_ ≈ 2.2 ×10^5^ M^-1^) direct transfer events. If dsDNA has an additional “foothold” of binding interactions on PRC2 not utilized by G4 RNA (Fig. 4), then under our model (Fig. 8b) it’s expected that dsDNA would be more resistant to displacement by an incoming RNA molecule and more efficient at displacing an existing RNA molecule.

### Implications for Other Chromatin Modifiers

Accumulating evidence suggests that RNA-binding activity is common among chromatin-associated proteins (35–39). As discussed above, our findings provide mechanistic evidence for a robust RNA-mediated regulatory function of one chromatin-modifying protein, PRC2. More importantly, however, they raise the possibility that this direct transfer mechanism might be general for RNA-mediated regulation. Historically, polynucleotide transfer mechanisms have been minimally studied and limited to oligomeric proteins (23–25, 27, 31). Our proposed mechanism (Fig. 8b) implicates dynamic protein-polynucleotide binding interactions as a key contributor to direct transfer. Under this model, the prevalence of oligomeric proteins in prior studies is logical, since these oligomers have multiple active sites that can dynamically bind ligand. Furthermore, our mechanism raises the question of how common direct transfer kinetics might be among nucleic acid binding proteins. In concurrent studies (companion manuscript) we interrogated the prevalence of direct transfer among numerous protein-ligand interactions, including a diverse range of nucleic acid binding proteins, and our findings suggest that direct transfer is quite common among other chromatin-associated proteins.

If direct transfer capability is indeed pervasive among chromatin-associated proteins, then our findings for PRC2 may explain why so many chromatin-associated proteins exhibit RNA-binding activity: intrinsic direct transfer capability allows for tunable RNA-mediated regulation. It’s important to distinguish the characteristics of competitive binding (PRC2-like) mediated direct transfer (Fig. 5) versus independent binding (YY1-like) mediated direct transfer (Fig. 6). In the former case, RNA can recruit protein under high RNA-nucleosome proximity conditions, but actively antagonize protein activity under low RNA-nucleosome proximity conditions. In the latter case, while RNA could indeed recruit proteins if there’s high RNA-nucleosome proximity, it would be unable to antagonize protein activity if RNA is predominantly in free solution.

Thus, it’s possible that chromatin-associated proteins’ have evolved PRC2-like versus YY1-like direct transfer in response to physiological pressures for tight-regulation versus efficient recruitment. Future *in vitro* and *in vivo* studies with a diversity of chromatin-associated proteins are warranted to interrogate the prevalence and nature of the direct transfer mechanism’s role(s) in RNA-mediated regulation of gene expression.

## MATERIALS & METHODS

### PRC2 Expression and Purification

According to prior methodology (33), we used pFastBac vectors encoding N-terminally MBP-tagged fusions of each of the four core PRC2 subunits (EZH2, SUZ12, EED, RBBP4), and either the embryonic (PRC2_5me_) or somatic (PRC2_5m_) isoform of AEBP2 (40), to prepare respective baculovirus stocks for co-infection of Sf9 cells. Then, according to prior methodology (14), cell paste containing expressed PRC2 was lysed, clarified, then purified by sequential amylose column chromatography, MBP-tag cleavage, heparin column chromatography, and size-exclusion column chromatography. The PRC2_5me_ protein was used for all reported experiments, except where otherwise indicated (see Table 1).

### Preparation of Polynucleotides

All oligos were ordered from IDT (Newark, NJ), and their sequences in IDT syntax are provided (Supp. Table 1). For dsDNA constructs, complementary oligos ordered from IDT were mixed at 5 µM (prey) or 300 µM (decoy) each in annealing buffer (50 mM TRIS pH 7.5 at 25°C, 200 mM NaCl), subjected to a thermocycler program (95°C for 10-min, 95→4°C at 0.5°C/min, hold at 4°C) for annealing, then annealing confirmed via Native-PAGE. Concentrations of all ligands were confirmed spectroscopically using manufacturer-provided extinction coefficients.

### Binding Buffers

All binding buffers (BB) contained 50 mM TRIS (pH 7.5 at 25°C), 2.5 mM MgCl_2_, 0.1 mM ZnCl_2_, 0.1 mg/mL BSA, 5% v/v glycerol, and 2 mM 2-mercaptoethanol, plus a variable concentration of KCl (10, 25, 100, or 200 mM). Subscript of each binding buffer indicates the concentration of KCl in milli-molarity (e.g., BB_25_ = 25 mM KCl).

### FP-Based K_d_ Determination

Pre-reaction mix was prepared with 5 nM prey molecule in respective binding buffer (see Binding Buffers), then dispensed in 36 µL volumes into the wells of a 384-well black microplate (Corning #3575). PRC2 was prepared at 10X the reported concentrations via serial dilution in binding buffer. Binding reactions were initiated by addition of 4 µL of PRC2 solution to the corresponding pre-reaction mix, then incubated 30 min at room temperature. Wells with binding buffer only were also included for blanking. Fluorescence polarization readings were then taken for 30 min in 30 s intervals with a TECAN Spark microplate reader (Excitation wavelength = 481 ± 20 nm, Emission wavelength = 526 ± 20 nm). Each experiment had 2 or 4 technical replicates per protein concentration (as indicated), and at least three independent experiments were performed per protein-polynucleotide combination.

Raw data were analyzed in R v4.1.1 with the FPalyze function (FPalyze v1.3.1 package; see Software, Data, and Materials Availability). Briefly, polarization versus time data were calculated for each reaction, the last 10 data points for each reaction averaged to generate an equilibrium polarization value, and then equilibrium polarization values were plotted as a function of protein concentration. Plot data were regressed with Eq. 2 to calculate K_d_^app^for the interaction.

### FP-Based Competitive Dissociation Experiments

Pre-reaction mix was prepared with 5 nM prey molecule and PRC2 ≥ 2xK_dP_^app^(at 25°C) in binding buffer (see Binding Buffer Compositions), then dispensed in 36 µL volumes into the wells of a 384-well black microplate (Corning #3575). Decoy was prepared at 10X the reported concentrations via serial dilution in binding buffer or carrier polynucleotide (Table 1) at a concentration equal to the highest decoy concentration. Pre-reaction mix and decoy dilutions were then incubated at the indicated temperature to attain thermal and binding equilibrium (4°C/90min or 25°C/30min).

Competitive dissociation reactions were initiated by addition of 4 µL of the respective decoy concentration to the corresponding pre-reaction mix, then fluorescence polarization readings were immediately (the delay between initiation of the first reactions and the first polarization reading was ∼90 s) taken at 25°C for 120 min in 30 s intervals with a TECAN Spark microplate reader (Ex = 481 ± 20 nm, Em = 526 ± 20 nm). Each experiment had 4 technical replicates per decoy concentration, and at least three independent experiments were performed per protein-polynucleotide combination unless otherwise indicated. All reported competition reactions used a PRC2 concentration of 100 nM.

Raw data were analyzed in R v4.1.1 with the FPalyze function (FPalyze v1.3.1 package). Briefly, polarization versus time data were calculated for each reaction, each reaction’s polarization data were normalized to the maximum and minimum polarization across all reactions, each normalized reaction was fit with an exponential dissociation function (Eq. 3.1) to determine k_off_^obs^ (Eq. 3.2), and k_off_^obs^values were plotted as a function of decoy concentration. Plotted data (with background k_off_ ^obs^subtracted to mitigate temperature effects on polarization) were regressed (the theoretical background for this approach is thoroughly covered in a separate manuscript) via Eq. 4.1 and then Eq. 4.2 with tuning parameters constrained to the Eq. 4.1 solutions, then the regression models compared with the Bayesian Information Criterion (41) (BIC). Rate constants (k_-1P_and/or k_θD_) were determined from the best-performing regression model. If minimum polarization was not reached during competition experiments (e.g., due to a weak competitor), then it was manually defined with minimum polarization data from corresponding binding curve data (see FP-Based K_d_ Determination).

### Ionic Strength Dependence

Binding curve data (Fig. 4a) were regressed as described to determine K_d_^app^ (see FP-based K_d_ Determination). Then, log_10_{K_d_^app^} versus log_10_{[KCl]} plots were regressed in R via stats:lm (package:function). Regression indicated m = 5.7×10^−4^ ± 0.34 (slope) and b = -8.5 ± 0.49 (intercept) for the G4 RNA data, and m = 1.4 ± 0.68 and b = -5.2 ± 1.1 for the dsDNA data, where values are the regressions’ estimate ± standard error.

### PRC2 Reaction Scheme Simulations

Reactions (Fig. 5a) were simulated and analyzed in R v4.1.1 with a custom script (see Software, Data, and Materials Availability). Briefly, [E_T_], [N_T_], [R_T_], K_d_, k_-1_, k_θ_, k_cat_, and α were user-provided. Then, other rate constants and initial conditions were calculated via Eq. 7, and the system of differential equations (Eq. 5) was solved by numerical integration. By default, k_-1_, k_θ_, and K_d_ values were taken directly from the Fig. 3 studies (Table 1; k_-1_ are from self-competitions), k_cat_ = 1 s^-1^, [N_T_] = 5 nM, and all other values are indicated. By exception, k_θ_ = 0 for the Supp. Fig. 6 studies, and k_-1R_ = 9 ×10^5^ s^-1^ and K_dR_ = 3 M for the Supp. Fig. 5 studies.

### Co-Binder Reaction Scheme Simulations

Reactions (Fig. 6a) were simulated and analyzed in R v4.1.1 with a custom script (see Software, Data, and Materials Availability). Briefly, [E_T_], [N_T_], [R_T_], K_d_, k_-1_, k_cat_, α, β, and δ were user-provided. Then, other rate constants and initial conditions were calculated via Eq. 7, and the system of differential equations (Eq. 6) was solved by numerical integration. Initial reaction rates (V_0_) were calculated as the average rate of change in [m_T_] during the first 10% of each reaction. By default, K_d_ values were estimated from Sigova et al. (34) (K_dR_ = 400 nM, K_dN_ = 200 nM), k_-1_ = K_d_ ×10^5^ s^-1^, k_cat_ = 10^−3^ s^-1^, [N_T_] = 50 nM, δ = 1, and all other values are indicated. By exception, δ_1_ = 0 for the Supp. Fig. 7 & 9c studies, δ_2_ = 100 for the Supp. Fig. 9d & 10 studies, and k_cat_ = 0 s^-1^ for the Supp. Fig. 9 studies.

### Single-Molecule Dynamics Simulations

Reactions (Fig. 7) were simulated and analyzed in R v4.1.1 with custom scripts (see Software, Data, and Materials Availability). Briefly, ‘nucleosome’ and ‘protein’ molecule coordinates were randomly scattered in a 3-dimensional simulation box with periodic boundary conditions, ‘RNA’ molecule coordinates were added linearly to both sides of each nucleosome in intervals of twice the molecular diameter (2x p_d_), nucleosome and RNA coordinates were checked for molecular clash (proximity ≤ p_d_), and coordinate generation was repeated if necessary. Then, molecular diffusion was approximated by ‘random walk’ changes in coordinates at each time-step, and inter-molecular binding was defined at the end of each time-step as a proximity of p_d_ or less between protein and nucleosome/RNA molecules. Molecules determined to be bound were set to a bound state, with their diffusion and additional binding capacity ablated, for a randomly sampled length of time based on the molecular pair’s dissociation rate constant (k_-1_).

Specifically, [E_T_], [N_T_], R_nN_ (RNA molecules per nucleosome), K_d_, t (reaction time), τ (time-step), and ζ (dimensions of cubic simulation box) were user-provided, D (diffusion coefficient), p_d_ (molecular diameter), and A_n_ (Avogadro’s number) were established constants, and D_µ_ (average diffusion distance per time-step), k_1_ (the association curves in our data suggest that our calculation for k_1_ may have underestimated the effective association rate produced by D in our simulations), and k_-1_ were calculated from other parameter values via Eq. 8.1-3. To generate initial conditions, numbers of nucleosome (N_n_) and protein (E_n_) molecules calculated by rounding Eq. 8.4 to the nearest integers, numbers of RNA molecules (R_n_) calculated via Eq. 8.5, initial nucleosome and protein cartesian coordinates sampled from a uniform distribution parameterized by [-ζ ÷2, ζ ÷2], RNA cartesian coordinates calculated by 2x p_d_-interval additions and subtractions to the nucleosome x-coordinates, an intermolecular distance of 2x p_d_ or greater confirmed between all nucleosome and RNA molecules, then coordinate assignment repeated if necessary. To simulate diffusion between time-steps for each protein molecule in an unbound state, spherical coordinates for direction were sampled from a uniform distribution parameterized by [0, 2π], the spherical coordinate for magnitude was sampled from an exponential distribution parameterized by D_µ-1_ (inverse of average diffusion distance per time-step), then spherical coordinates converted to cartesian coordinates and added to the existing coordinate values. Molecules that diffused past a ‘wall’ in the defined simulation box during each time-step were moved a proportionate distance into the simulation box from the opposite ‘wall’ (periodic boundary conditions). To determine binding states after initial conditions and each diffusion step, the intermolecular distance to every protein molecule was calculated sequentially for every nucleosome/RNA molecule, the most proximal protein molecule with an intermolecular radius of p_d_ or less was identified (if any), a binding state value of zero (unbound) confirmed for the protein and nucleosome/RNA molecules, a residence time sampled from an exponential distribution parameterized by k_-1_ and rounded to the corresponding integer number of time-steps, and the time-step number set as the new binding state value. Zero binding states indicate unbound molecules, non-zero binding states indicate bound molecules, and non-zero binding state values drop by 1 at the end of each time-step (after diffusion and binding state updates).

Nucleosome/RNA occupancy was calculated as the fraction of nucleosome/RNA molecules with non-zero binding states, nucleosome-protein proximity was calculated as the average intermolecular distance between every nucleosome molecule and their closest R_nN_ ÷2 unbound protein molecules, and effective molarity was calculated via Eq. 8.6. By default, t = 50 ms, τ = 10 ns, ζ = 1 µm, K_dN_ = 500 nM, K_dR_ is indicated (Fig. 7), [E_T_] = 300 nM, [N_T_] = 15 nM, R_nN_ = 8, D = 100 µm^2^ s^-1^, p_d_ = 5 nm, and A_n_ = 6.022 ×10^23^ mol^-1^.

### Diagram, Reaction Scheme, and Figure Generation

Diagrams were prepared with BioRender, reaction schemes were prepared with ChemDraw v21.0.0 (Perkin Elmer), tables were prepared with Word (Microsoft), graphs were prepared with R v4.1.1, protein structures were prepared in PyMOL v2.5.2 (Schrodinger), and figures were assembled in PowerPoint (Microsoft).

### Software, Data, and Materials Availability

GitHub hosts the FPalyze (github.com/whemphil/FPalyze) R package. The custom scripts referenced in these methods are available on GitHub (github.com/whemphil/PRC2_Direct-Transfer_Manuscript). Requests for other data and materials should be directed to T.R.C.

### Equations

For Eq. 1.1-5, rate constants are defined in Fig. 2 – step 3, E is protein (PRC2), P is prey, D is decoy, conjugations of these reactants are complexes, equations give rates of change for indicated reactants as a function of time (t), and bracketed terms indicate concentrations. For Eq. 1.6-10, apply Eq. 1.1-5 notation.

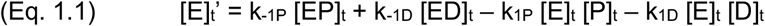

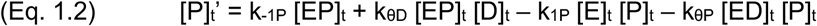

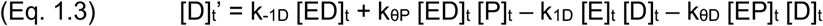

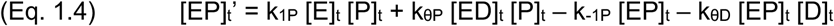

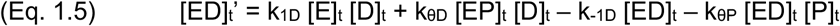

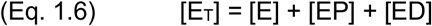

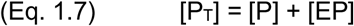

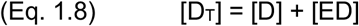

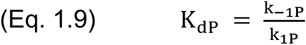

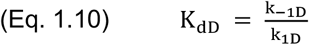

For Eq. 2, P_E_ is polarization at equilibrium for a given [E_T_], P_max_ is the maximum polarization, P_min_ is the minimum polarization, [E_T_] is the total protein concentration, and K_dapp_ is the apparent dissociation constant.

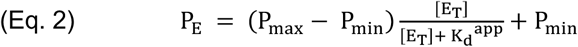

For Eq. 3.1-2, N_t_ is relative polarization at a given time (t), N_min_ is the minimum relative polarization, λ is the decay rate constant, and k_off_^obs^is the observed dissociation rate.

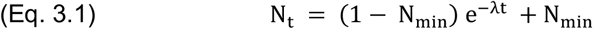

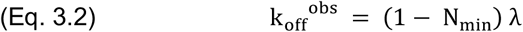

For Eq. 4.1-2, apply Eq. 1 notation, and α and β are arbitrary tuning parameters. These equations are derived in concurrent studies (separate manuscript).

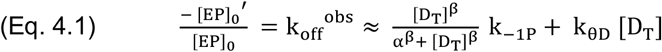

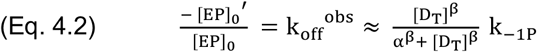

For Eq. 5.1-7, terms are defined in Fig. 5a, equations give rates of change for indicated reactants as a function of time (t), and bracketed terms indicate concentrations. For Eq. 5.8-11, apply Eq. 5.1-7 notation.

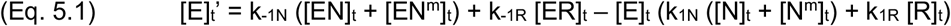

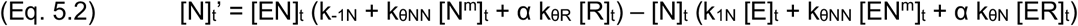

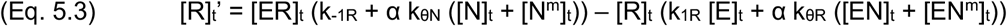

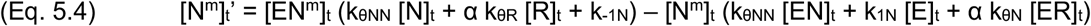

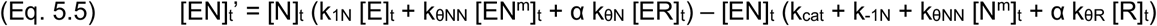

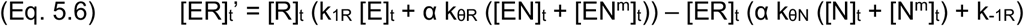

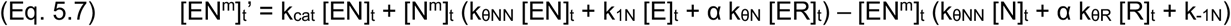

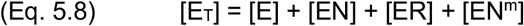

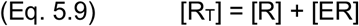

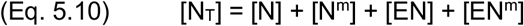

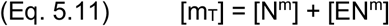

For Eq. 6.1-9, terms are defined in Fig. 6a, equations give rates of change for indicated reactants as a function of time (t), and bracketed terms indicate concentrations. For Eq. 6.10-13, apply Eq. 6.1-9 notation.

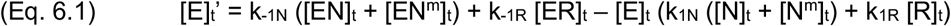

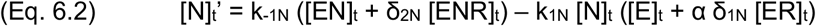

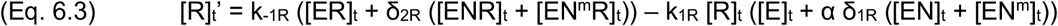

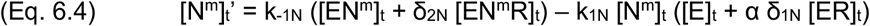

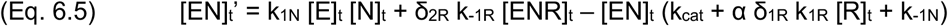

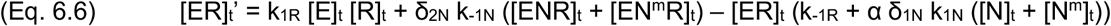

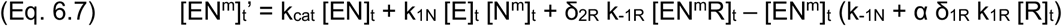

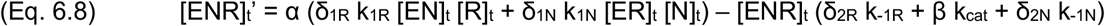

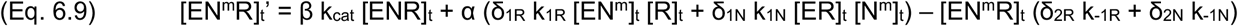

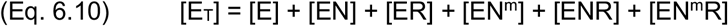

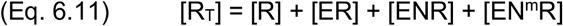

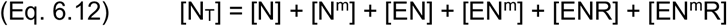

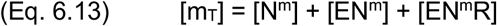

For Eq. 7.1-5, apply Eq. 5 notation. For Eq. 7.5b, apply Eq. 6 notation.

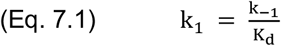

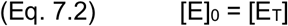

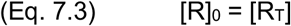

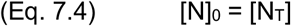

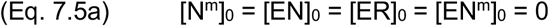

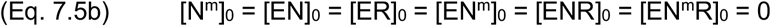

For Eq. 8.1-5, D_µ_ is the average diffusion distance in a time-step interval, τ is the time-step interval, D is the diffusion coefficient, X_n_ is the number of nucleosome/RNA molecules, A_n_ is Avogadro’s number, [X_T_] is the total concentration of nucleosome/RNA, ζ is the dimension length of a cubic simulation box, R_n_ is the number of RNA molecules, N_n_ is the number of nucleosome molecules, R_nN_ is the number of RNA molecules per nucleosome, M_ε_ is effective molarity, p_d_ is molecular radius, E_ζ_ ^app^ is the number of unbound protein molecules in the simulation box, and E_i_ is the number of unbound protein molecules within a 10x p_d_ radius of the i^th^ nucleosome molecule.

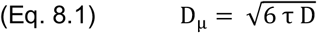

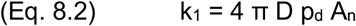

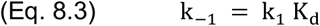

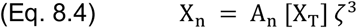

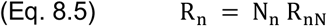

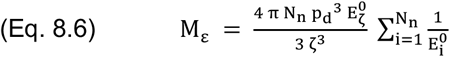

## ACKNOWLEDGEMENTS

W.O.H. was supported by the National Institutes of Health (F32-GM147934). T.R.C. is an investigator of the Howard Hughes Medical Institute.

We’d like to thank Olke Uhlenbeck, Deborah Wuttke, Halley Steiner, and members of the Cech lab (University of Colorado Boulder), for stimulating discussion and feedback concerning these studies.

---

**Supplemental Table 1.**
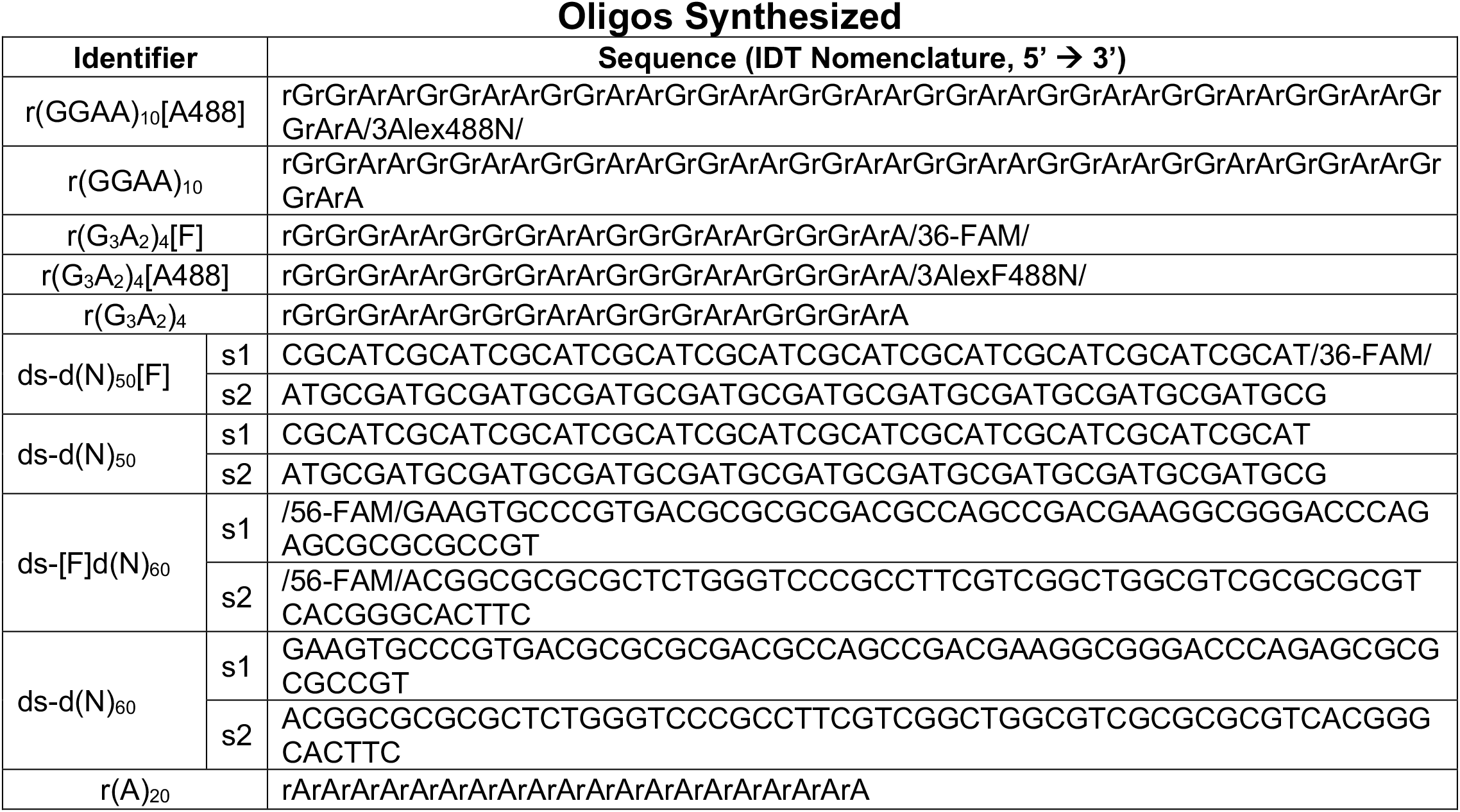
Identities of Synthetic Polynucleotide Species. The nomenclature used for ordering oligos from IDT (Sequence) is provided for all oligos named (Identifier) in these studies.

**Supplemental Figure 1.**
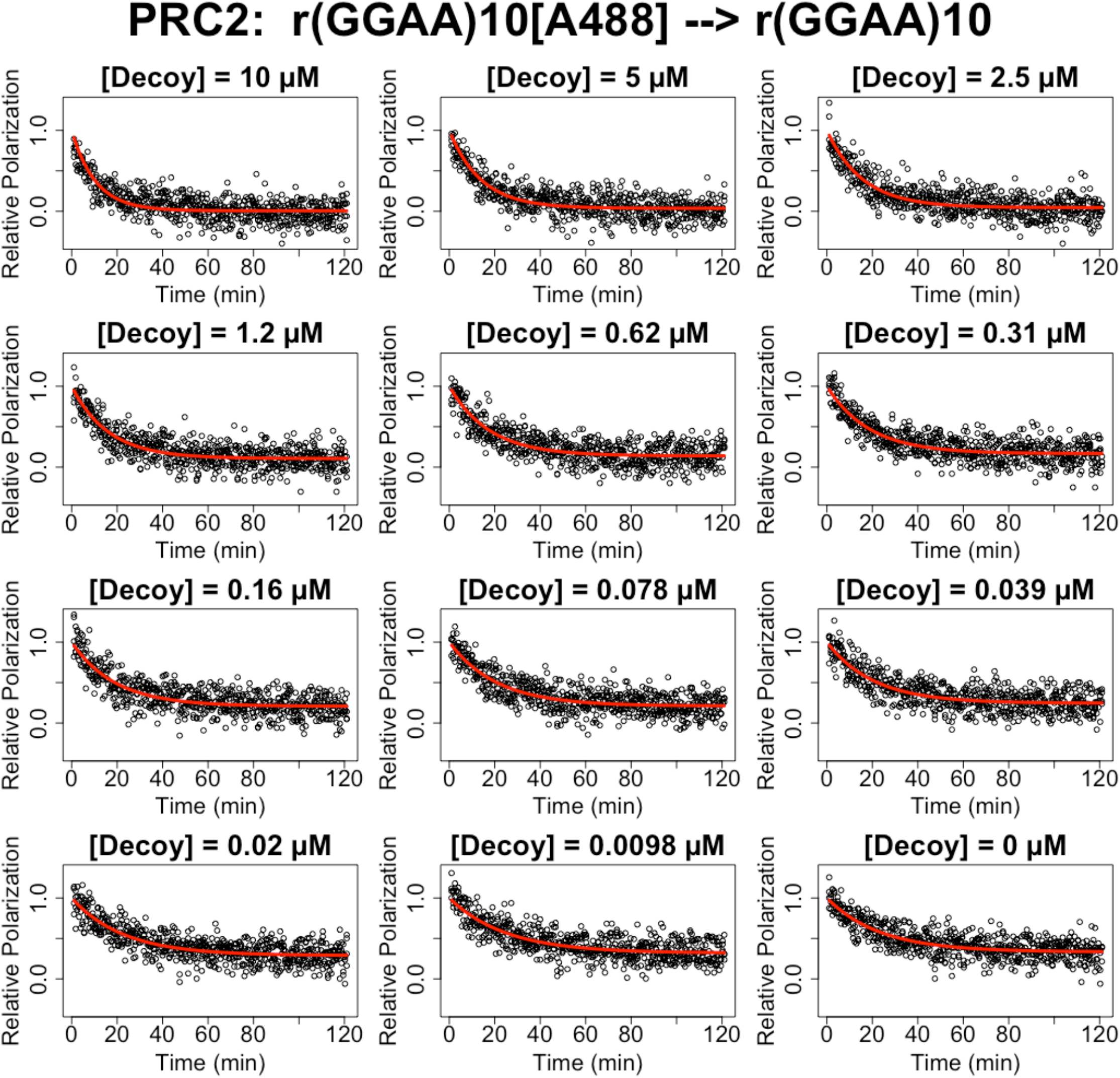
Raw Data for Figure 1. Fluorescence polarization-based competitive dissociation experiments (Fig. 2) were performed as described to replicate the original Wang et al and Long et al experiments over a range of competitor (‘decoy’) RNA concentrations. Raw data is from the same single representative experiment (of n=3) as Fig. 1, with four technical replicates. Reaction data for each competitor concentration is regressed with an exponential dissociation equation (Eq. 3.1), and the fit lines are shown (red lines).

**Supplemental Figure 2.**
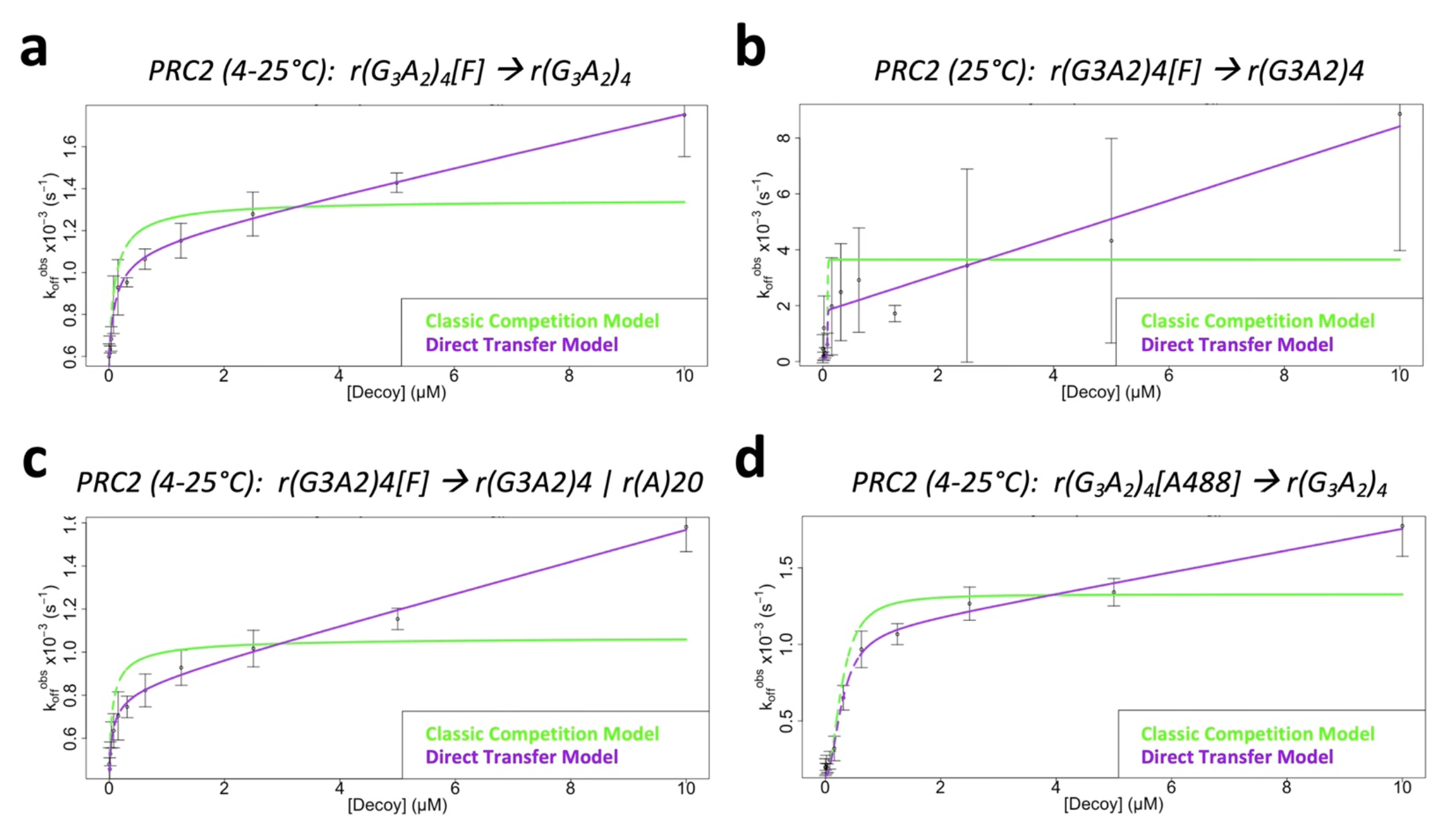
A Simpler G4 RNA Exhibits Direct Transfer Kinetics Independent of Temperature, Polynucleotide Concentration, and Fluorescent Label. FPCD experiments (Fig. 2) were performed (buffer = BB_25_) as described for a simplified G-quad RNA as prey and decoy. Data are from representative experiments (of n=3), where error bars indicate mean ± SD for four technical replicates. **[a]** Standard FPCD experiment. **[b]** FPCD experiment at constant 25°C – control for variable temperature artifacts. **[c]** FPCD experiment with nonbinding carrier RNA to keep total RNA concentration constant – control for nonspecific RNA concentration-dependent artifacts. **[d]** Standard FPCD experiment with different fluorophore.

**Supplemental Figure 3.**
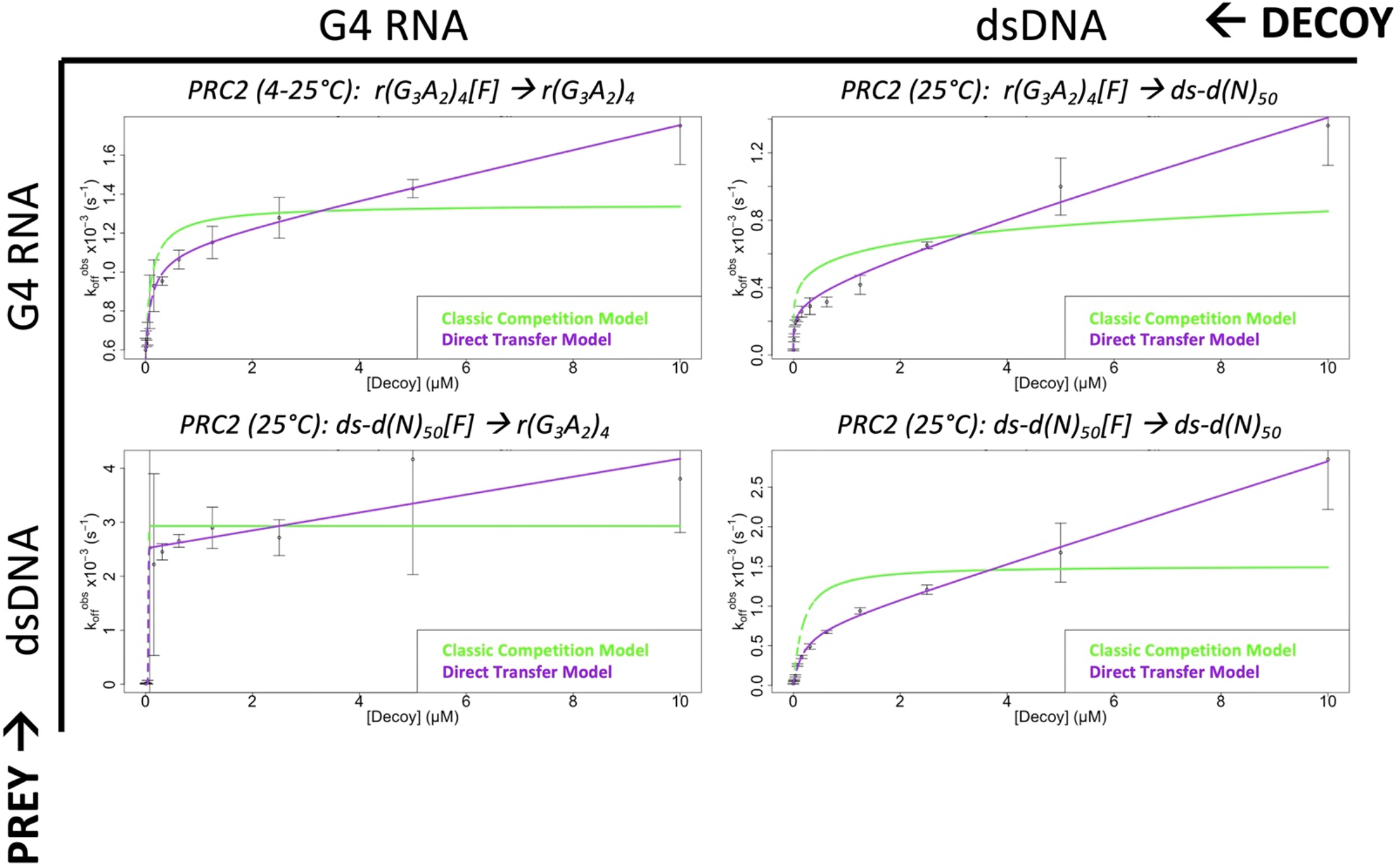
PRC2 Exhibits Direct Transfer Kinetics for G4 RNA and Another dsDNA. FPCD experiments (Fig. 2) were performed (buffer = BB_10_) as described for every prey-decoy combination of a G-quad RNA and 50-bp dsDNA. Data are from representative experiments (of n=3), where error bars indicate mean ± SD for four technical replicates.

**Supplemental Figure 4.**
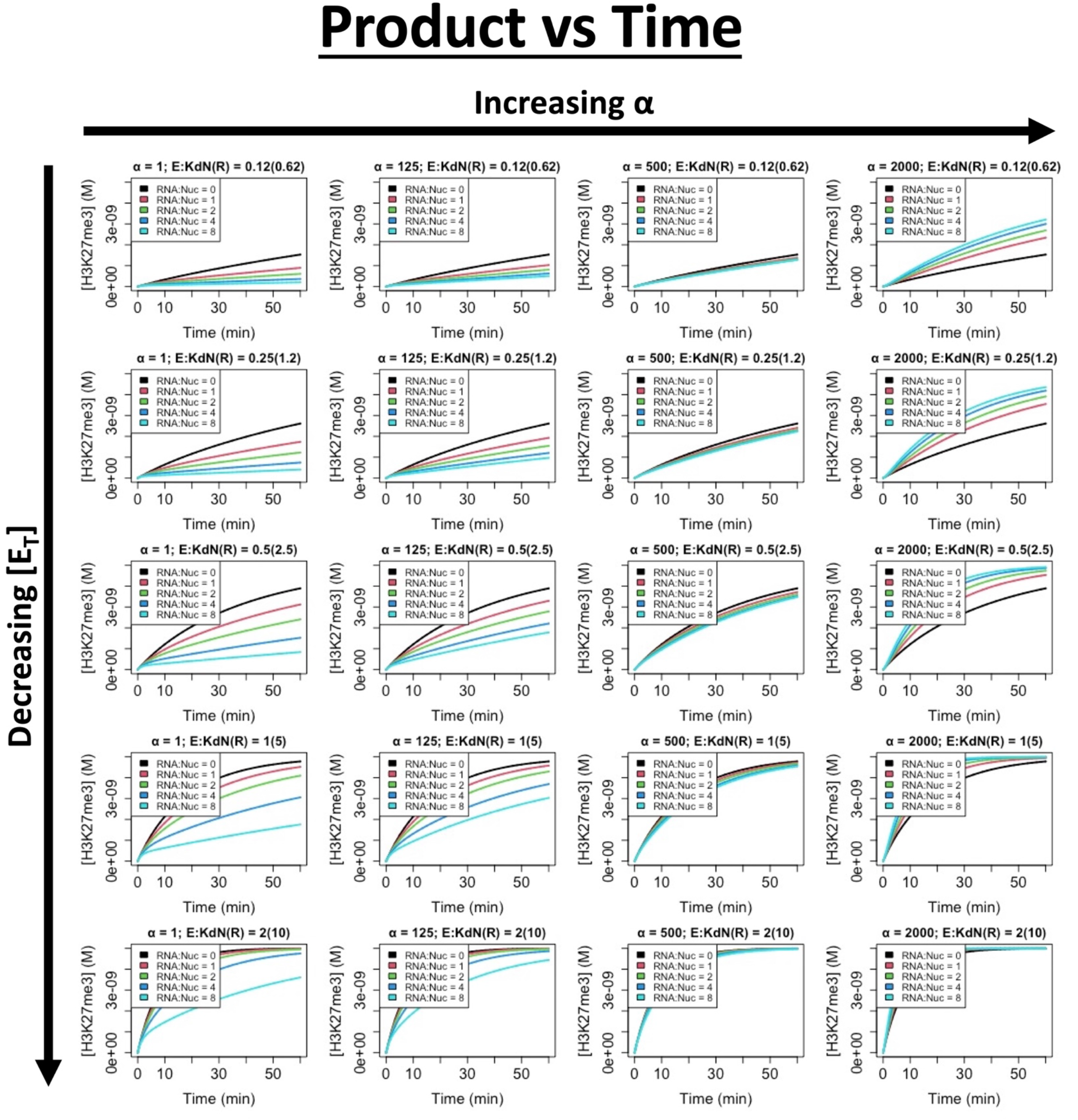
Full Data Set for Figure 5. Empirically determined rate constants (Table 1) were used to simulate PRC2 HMTase activity for a range of effective molarities (α), PRC2 concentrations (E), and RNA-nucleosome molar ratios (RNA:Nuc).

**Supplemental Figure 5.**
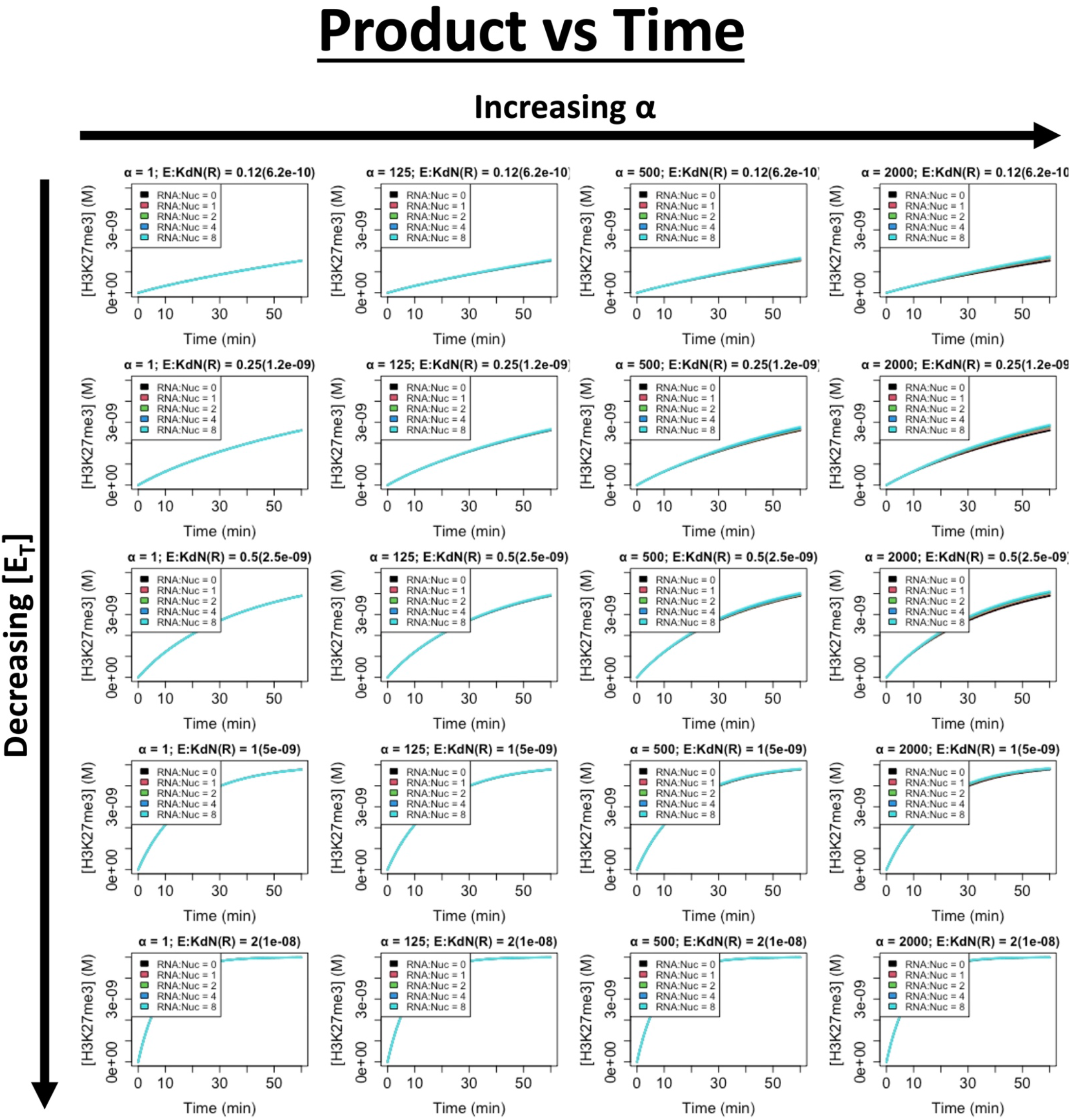
Unstable RNA Binding Ablates RNA-Dependent Effects on PRC2’s HMTase Activity. Supp. Fig. 4 simulations were altered to make RNA binding unstable (k_-1R_ = 9×10^5^ s^-1^, K_dR_ = 3 M).

**Supplemental Figure 6.**
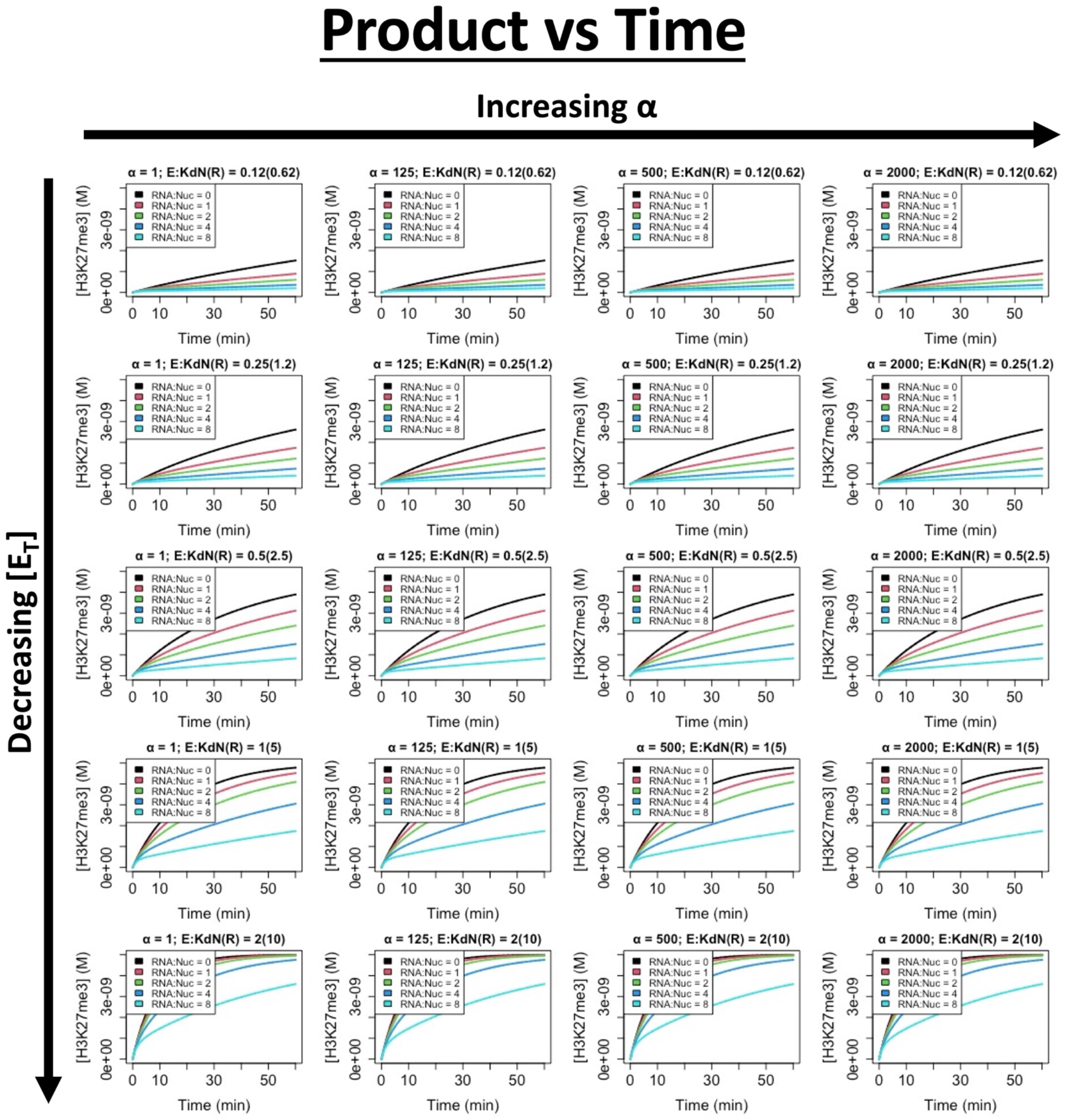
PRC2’s Proposed RNA-Dependent HMTase Boost is Dependent on Direct Transfer. Supp. Fig. 4 simulations were altered to eliminate direct transfer (k_θ_ = 0).

**Supplemental Figure 7.**
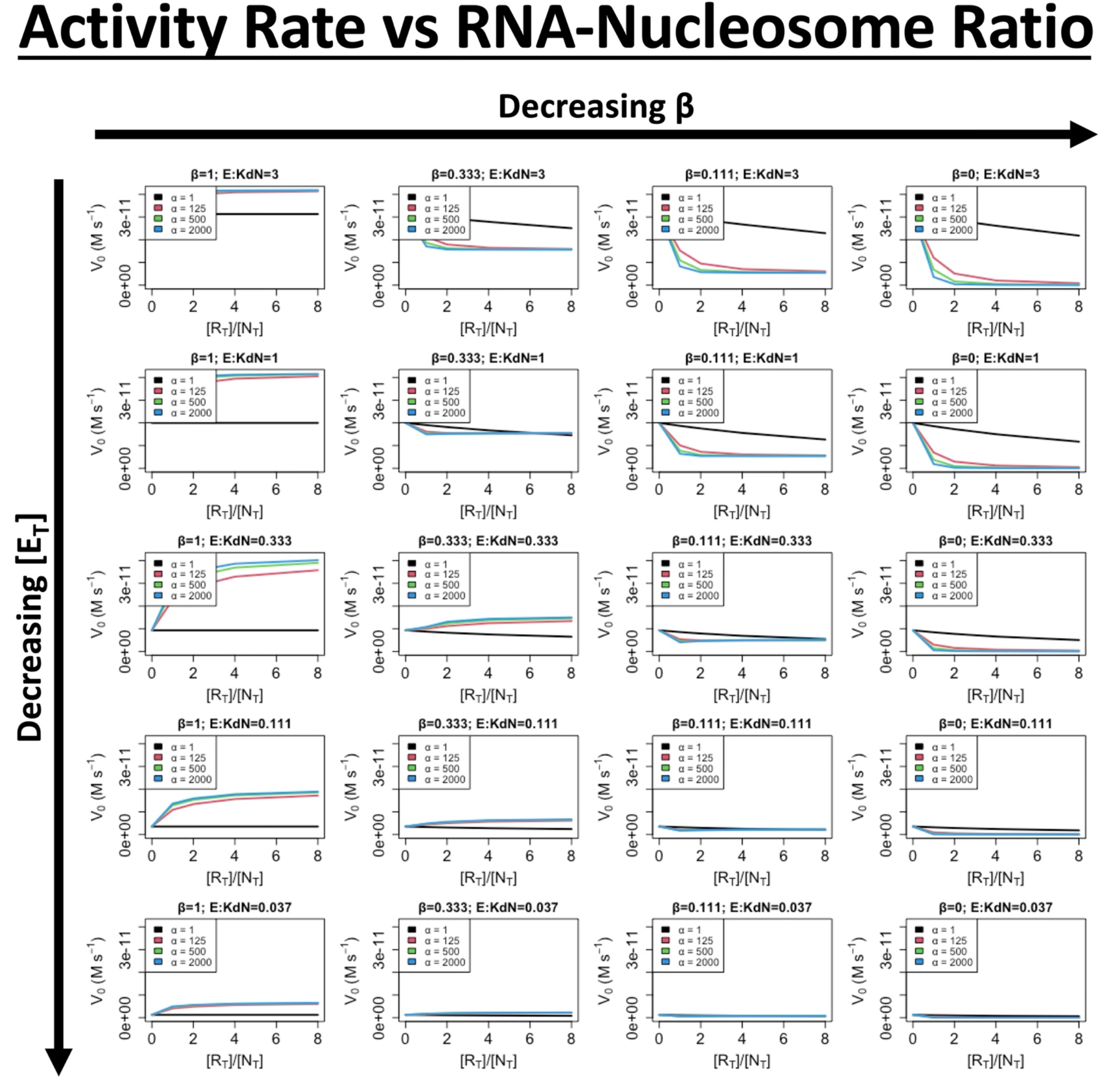
RNA-Dependent Activity Boosting for Co-Binders is Ablated by Minor RNA-mediated Catalytic Suppression. The Fig. 6 simulations were expanded to calculate initial reaction velocities (V_0_) for a range of effective molarities (α), levels of RNA-mediated suppression of catalysis (β), protein concentrations (E), and RNA-nucleosome molar ratios ([R_T_]/[N_T_]).

**Supplemental Figure 8.**
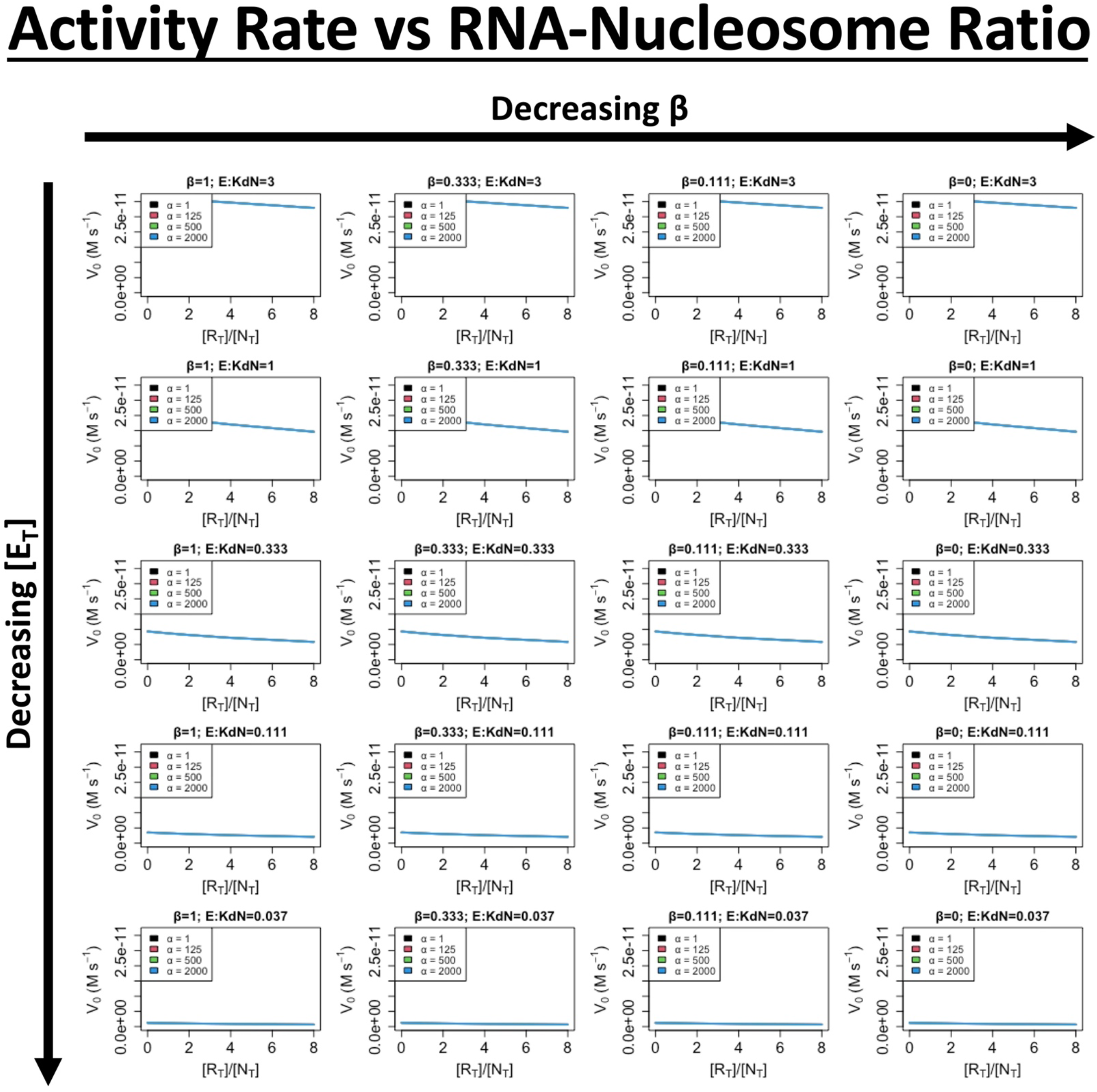
RNA-mediated Activity Boosting for Co-Binders is Dependent on Ternary Complex Formation. The Supp. Fig. 7 simulations were altered to prevent ternary complex formation (δ_1_ = 0).

**Supplemental Figure 9.**
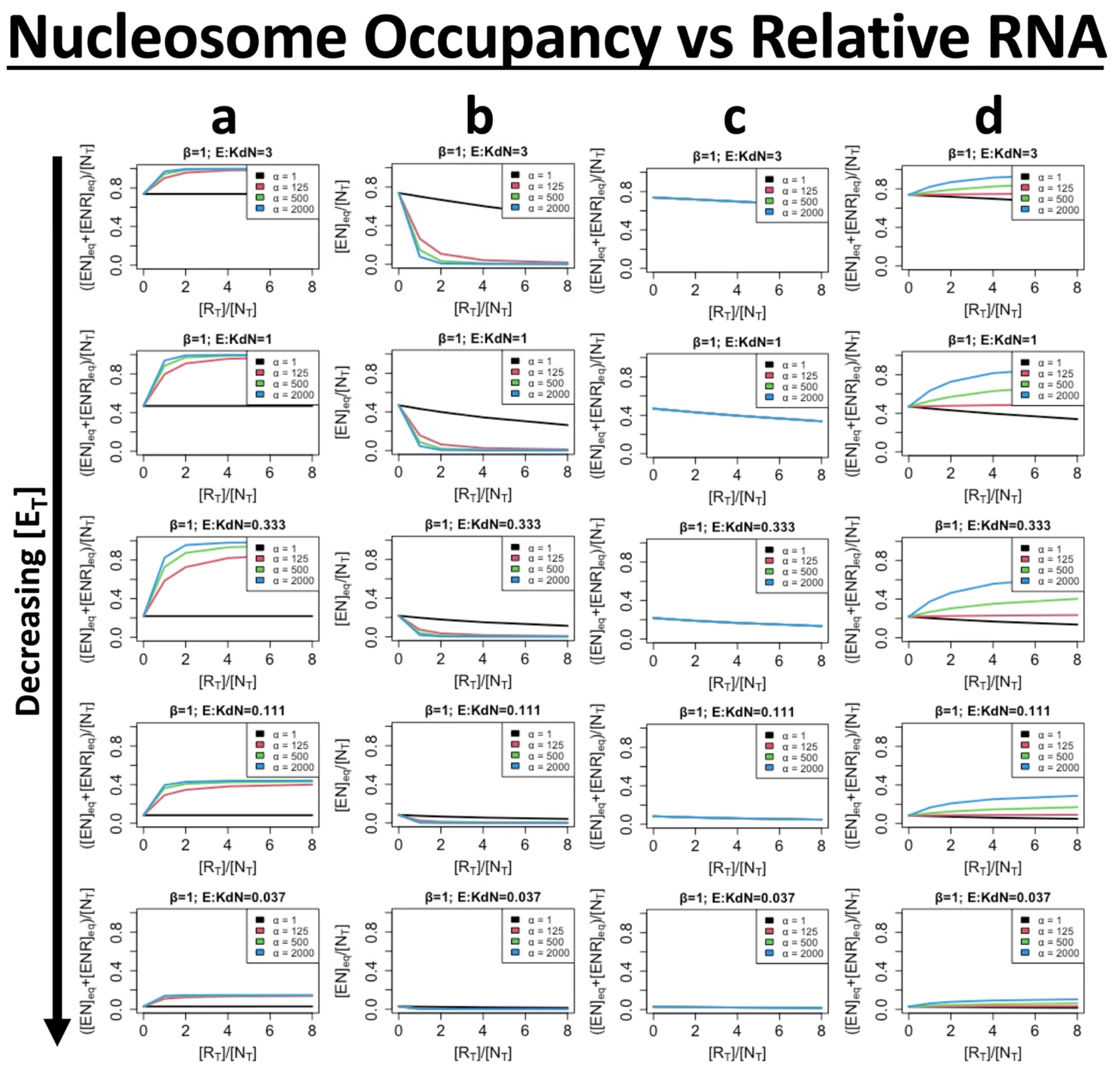
Trends in RNA-mediated Function for Non-Catalytic Co-Binders Resemble Trends for HMTase-Active Co-Binders. **[a]** *Co-Binding Boosts Nucleosome Occupancy*. The Fig. 6 reaction parameters were modified to disallow HMTase activity (k_cat_ = 0), then simulations performed to calculate equilibrium (eq) nucleosome occupancy for a range of effective molarities (α) and protein concentrations (E). The y-axis is the fraction of nucleosomes occupied by protein, either with or without the RNA remaining bound. The x-axis is the RNA-nucleosome molar ratio ([R_T_]/[N_T_]). **[b]** *RNA-mediated Suppression of Function During Nucleosome Occupancy Creates an Antagonistic RNA-Nucleosome Activity Relationship*. Panel (a) data are re-plotted with RNA-nucleosome co-occupancy excluded. **[c]** *RNA-mediated Nucleosome Occupancy Boosts are Dependent on Ternary Complex Formation*. The panel (a) simulations were altered to prevent ternary complex formation (δ_1_ = 0). **[d]** *An Unstable Ternary Complex is Sufficient for RNA-mediated Nucleosome Occupancy Boosting*. The panel (a) simulations were altered to drastically accelerate ternary complex dissociation (δ_2_ = 100).

**Supplemental Figure 10.**
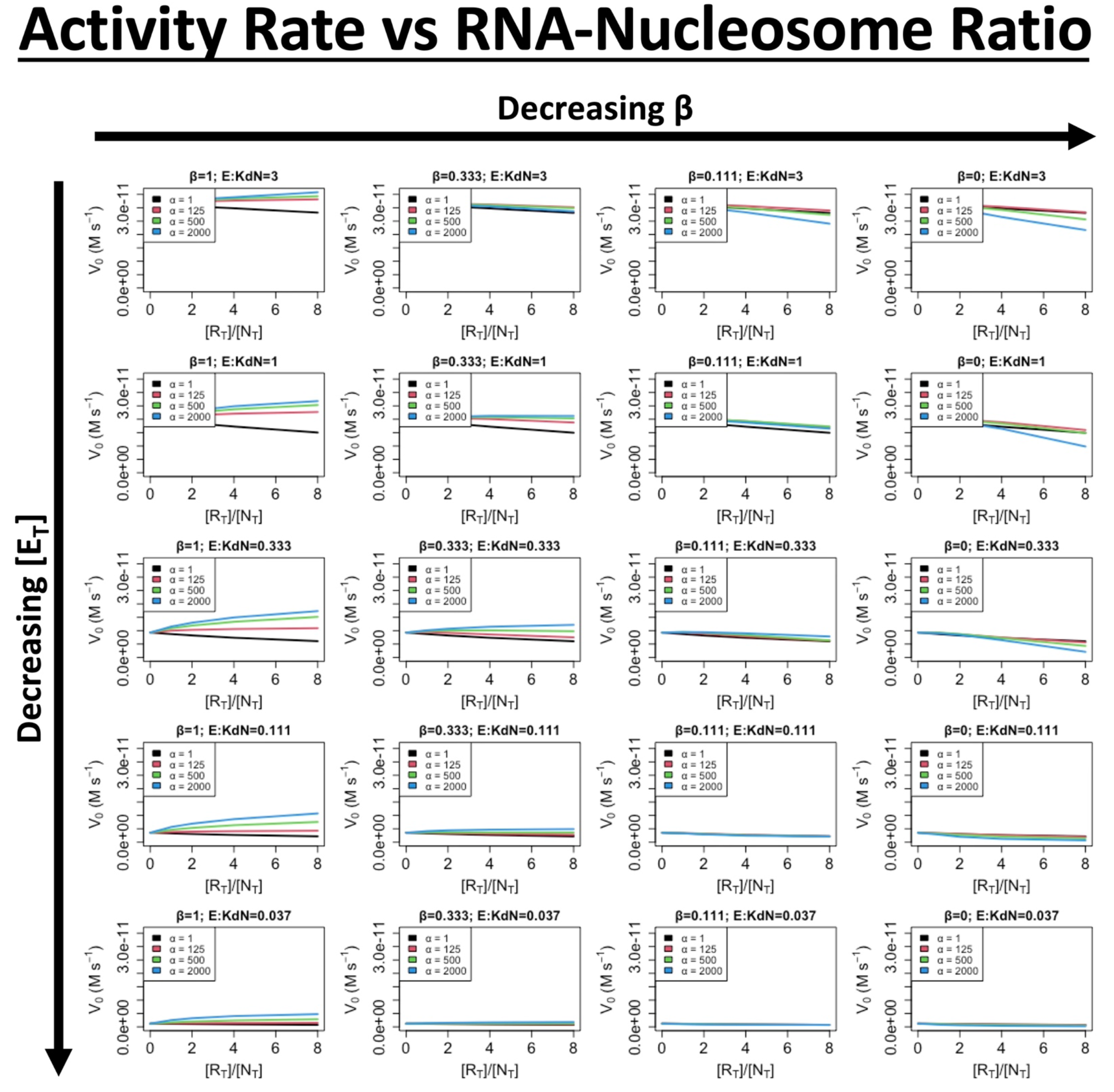
An Unstable Ternary Complex is Sufficient for Some RNA-mediated Activity Boosting for Co-Binders. The Supp. Fig. 7 simulations were altered to drastically accelerate ternary complex dissociation (δ_2_ = 100).

